# Disruption of The Spinal Cord-Gut Axis Alters Gut Microbial Dynamics and Carbohydrate Cross-feeding

**DOI:** 10.1101/2025.05.27.656368

**Authors:** Mohamed Mohssen, Ahmed A. Zayed, Kristina A. Kigerl, Jingjie Du, Garrett J. Smith, Jan M. Schwab, Matthew B. Sullivan, Phillip G. Popovich

**Affiliations:** The Interdisciplinary Biophysics Graduate Program, The Ohio State University, Columbus, Ohio 43210, USA; Department of Microbiology, The Ohio State University, Columbus, Ohio 43210, USA; Center of Microbiome Science, The Ohio State University, Columbus, Ohio 43210, USA; EMERGE Biology Integration Institute, The Ohio State University, Columbus, Ohio 43210, USA; Department of Neuroscience, The Ohio State University Wexner Medical Center, Columbus, OH, 43210, USA; Belford Center for Spinal Cord Injury, The Ohio State University Wexner Medical Center, Columbus, OH, 43210, USA; Department of Civil, Environmental and Geodetic Engineering, The Ohio State University, Columbus, OH 43210, USA

## Abstract

The spinal cord, a nexus for brain-body crosstalk, controls gut physiology and microbial homeostasis, but the underlying mechanisms remain unclear. Using genome-resolved longitudinal metagenomics in male and female C57BL/6 mice before and up to 6 months after disrupting the spinal cord-gut axis, we reconstructed over 6,500 microbial draft genomes. This “Mouse B6 Gut Catalog” improved or doubled species- and strain-level representation in other published catalogs. Impaired spinal cord-gut crosstalk induced persistent, sex-, time- and lesion-specific alterations in community composition, marked by a consistent loss of *Lactobacillus johnsonii*. Feeding this key bacterium to mice with a clinically relevant spinal cord injury improved host health. Genome-resolved, community-contextualized metabolic profiling revealed that shifts in carbohydrate-mediated microbe-microbe interactions explain the reduction of *L. johnsonii*. These findings identify carbohydrate metabolism as a keystone mechanism shaping gut microbiota and emphasize that mammalian health and gut ecosystem function depend on a functional spinal cord-gut axis. Additionally, these data improve murine microbiome catalogs and demonstrate that metagenome-informed microbial interventions can improve host health and likely mitigate long-term dysbiosis.

## INTRODUCTION

Mammals harbor a “second genome” composed of a complex microbial community residing in, on, and around their bodies. This microbial ecosystem, particularly prominent in the gut, assists the host in disease protection and influences the efficacy of drug therapies against illnesses like cancer^1^. These microbial beneficial roles depend on host control systems that monitor and regulate the microbiome^2^. One such system is the bi-directional signaling pathway connecting gut and brain functions^3,4^. Within this microbiota-gut-brain axis, the vagus (parasympathetic) nerve is often implicated as a solitary controller of host-microbiota mutualism^5,6^. However, the spinal sympathetic (fight-or-flight) nervous system also controls the gut, although its effects on the gut microbiota are rarely considered.

The spinal cord is the “hub” integrating nerve impulses before they influence the brain and body. Everyday stimuli, such as eating or bowel and bladder filling, activate spinal sympathetic nerves, affecting gut motility, mucus secretion, intestinal blood flow, and immune function^7^. Each of these gut physiological features can influence gut microbiota^8,9^. Even organ-specific signals directly transmitted to the brain via the vagus nerve, will directly^10,11^ and indirectly^12,13^ elicit spinal sympathetic reflexes that influence peripheral organs, including the gut^14^. Thus, to accurately assess gut microbiota stability in health and disease, a more holistic understanding of the “microbiota-gut-brain-spinal cord axis” is crucial.

Gut dysbiosis—imbalances in microbiome composition—is associated with the onset and severity of several health conditions, including central nervous system (CNS) diseases^15^. Accordingly, modifying gut microbiota is a focal point for therapeutic intervention. Despite wide use, gene-marker-based approaches like 16S rRNA gene sequencing^16–18^ face limitations, including primer bias, reliance on reference genomes, and restricted functional representation of niche-defining traits, which evolve faster than core traits and drive ecological interactions^19–21^. Genome-resolved metagenomics overcomes these limitations, enhancing taxonomic resolution and constructing genomic potential of ecosystem-specific taxa^22,23^. Genome-resolved metagenomics has been applied to the spinal cord-gut axis only once, in our prior proof-of-concept murine study limited by single timepoint and sex sampling^24^. Here, we longitudinally sample both sexes to create and curate a large, high-resolution murine microbiome reference genome catalog, then use it to evaluate the spinal cord control over the gut ecosystem, establishing a foundational understanding of how microbiota influence mammalian health. These advances deconstruct conventional thinking about gut-brain communication, showing that disruption to the spinal cord-gut axis causes sex- and time-dependent microbiota changes, altering microbial carbohydrate metabolism with implications for host health.

## RESULTS AND DISCUSSION

### Expanding murine microbial reference genomes

We first sought to build a more comprehensive murine microbial reference genomes catalog using an expanded experimental design, which is summarized as follows (details under “*Animals and spinal cord injury”* in **Methods**). First, microbiomes were compared from three groups of mice: (i) control mice receiving a sham-lesion to the spinal cord (Laminectomy or Lam; surgical removal of vertebral body but no spinal lesion), (ii) mice that received a complete lesion in low-thoracic (T10) spinal cord, and (iii) mice that received a complete lesion in high-thoracic (T4) spinal cord. Complete lesions at these spinal levels partially (T10) or completely (T4) abolish supraspinal control over spinal sympathetic neurons that control the gut^25,26^, effectively titrating the magnitude of “spinal cord-gut axis” dysfunction. Second, to enhance the longitudinal resolution of spinal cord-dependent changes (i.e., acute and chronic) on gut microbiota dynamics, samples were collected across six time points for each mouse. Third, to improve translational relevance and statistical power, we included female and male mice, and for each sex and spinal lesion level, we doubled the number of mice sampled at each time point.

Overall, these efforts resulted in 333 bulk metagenomes (294 fecal and 39 cecal) collected from 29 male and 30 female C57BL/6 mice at six time points spanning six months (**Fig. 1A**) – a considerable increase on our prior sampling^24^ of five female samples per experimental group (Lam, T4 and T10 spinal lesions) at a single time point. The overall physiological state of each mouse in the three groups was assessed after six months using a standardized scale of neurological recovery for mice (i.e., the Basso Mouse Scale (BMS) for Locomotion^27^; hereafter referred to as “BMS score” (**Fig. S1**). This expanded experimental design was further augmented by enhancing metagenomic workflow at several key steps, including cross-assembly per mouse, ensemble binning, and bin refinement (see **Methods** and **Fig. S1A**).

**Fig. 1.**
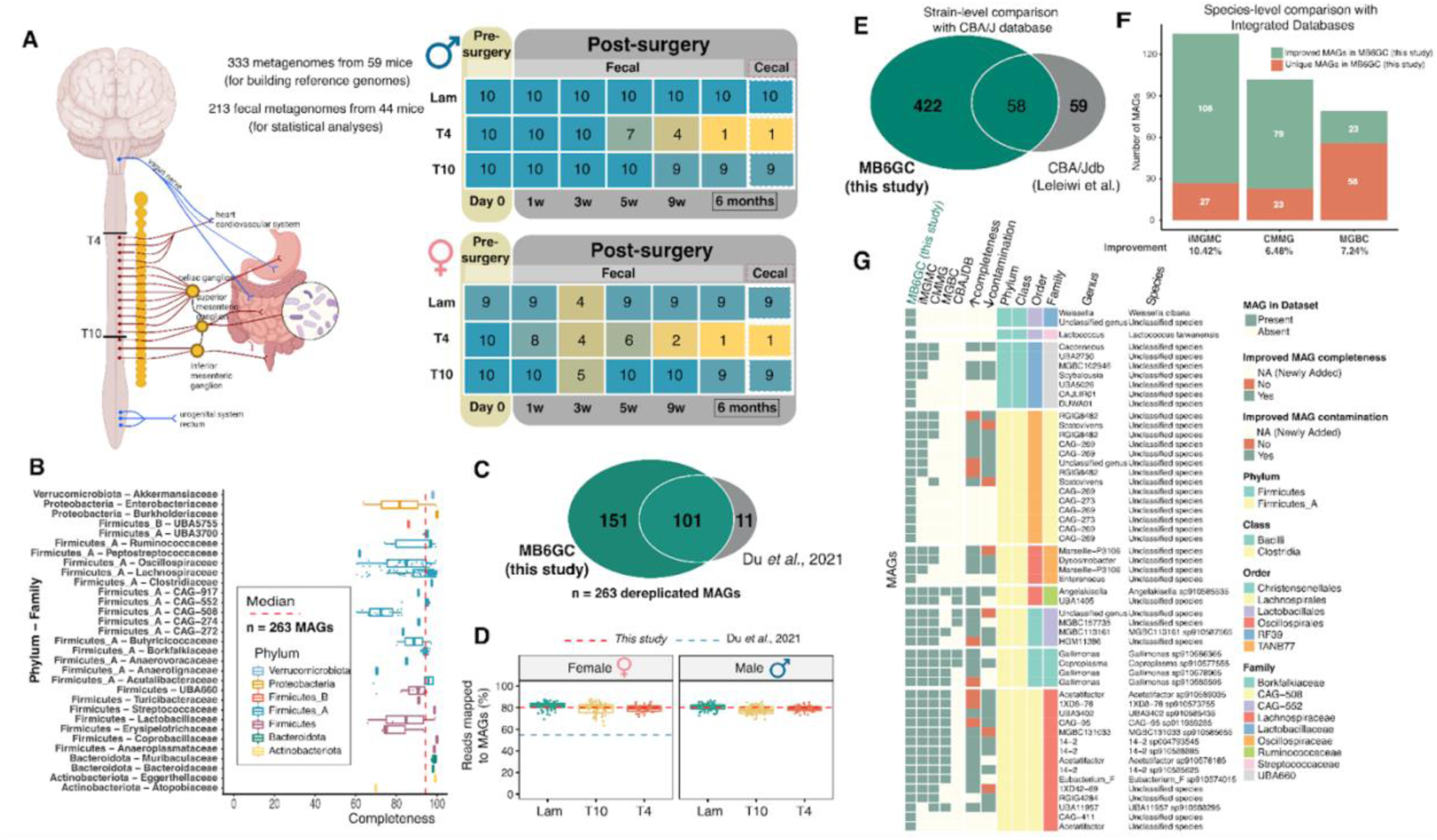
Study design, characterization of recovered MAGs and comparative analysis of Mouse B6 Gut Catalog (MB6GC) against other published mouse gut microbiome catalogs. **(A)** Schematic illustrates the relative location of spinal lesions and depicts the spinal sympathetic and vagal parasympathetic innervation to the gut. Tables (middle) describe the number of samples in this study, categorized by sex, spinal lesion level (Lam, T10, T4), and time post-lesioning. Note that autonomic complications in mice with lesions at spinal level T4 significantly reduce lifespan, yielding fewer T4 SCI mice, regardless of sex, beyond 7 weeks. (**B**) The completeness of the de-replicated MAGs (n=263; de-replicated at 95% ANI from 6,635 MAGs) is shown together with the phyla and families they span. New representative taxa include 2 phyla (Verrucomicrobiota and Proteobacteria), 6 families (Akkermansiaceae, Burkholderiaceae, Atopobiaceae, CAG-552, Enterobacteriaceae, CAG-272), and 41 genera (see GTDB data under Zenodo DOI). The median completeness across all MAGs is shown with a vertical dotted red line at 94.35%. Contamination levels are <10% for all de-replicated MAGs. (**C**) Venn diagram comparing the number of de-replicated MAGs recovered in this study against those from our prior pilot study^24^. (**D**) The fraction of reads recruited to the de-replicated set of MAGs, where the mean (red dotted line) is ∼80% across sexes and spinal injury levels (also see SingleM supporting data in **Fig. S1B**). The mean for^24^ (blue dotted line) was only available for female mice. The mean number of reads mapped to MAGs, as evaluated by the Wilcoxon test, was significantly improved in the current study (*p*-value= 6.11e-11). Schematic created with BioRender. **(E)** Venn diagram showing that at the MAG strain level (99% ANI dereplication), MAGs from MB6GC (this study; C57BL/6 mice) are largely mouse-strain specific (see **Methods**), with minimal overlap with the CBA/J MAG catalog. (**F**) A stacked bar chart shows that at the MAG species level (95% ANI dereplication), MAGs from MB6GC improve upon each of the previously published catalogs of mouse-gut reference genomes, including the Comprehensive Mouse Microbiota Genome Catalog (CMMG), the Mouse Gastrointestinal Bacteria Catalogue (MGBC), and the Integrated Mouse Gut Metagenomic Catalog (iMGMC). (**G**) A taxonomically resolved heatmap shows that 53 MAGs from MB6GC were either novel (n=15) or were the best representative genomes of their species (n=38) compared to other catalogs. MB6GC’s improvements in genome completeness and contamination over all catalogs are shown.

The increased sampling and updated analytics drastically expanded microbiota representation. In total, recovered genomic representation increased with 6,635 bacterial metagenome-assembled genomes or MAGs (>50% completeness and <10% contamination), that when combined with 112 previously generated MAGs^24^ led to 263 de-replicated MAGs (at >95% average nucleotide identity, or ANI) spanning 7 phyla and 31 families (**Fig. 1B**), which were used for all ecological and metabolic analyses. Of the 263 de-replicated MAGs, 238 were either above-medium (n = 76; >70% complete, <10% contamination) or high (n = 162; >90% complete, <5% contamination) quality (see **Methods**) – more than doubling MAG-based genomic representation from our prior pilot study^24^ (**Fig. 1C**). Taxonomically, this larger MAG collection captured additional representative taxa above the species level (**Fig. 1B**). Furthermore, 20.5% (n=54/263) of newly identified MAGs are likely novel species-level taxa as they lacked 95% Average Nucleotide Identity (ANI)-level representation, even against the ∼62k bacterial species clusters from the many environments represented in the Genome Taxonomy Database (GTDB^28^; Release 207). Ecologically, this updated MAG collection significantly improved the representation of murine gut microbiota as the percentage of sequencing reads that align to de-replicated MAGs increased from a mean of 55%^26^ to ∼80% (**Fig. 1D**). Further, this improved MAG-based read-mapping finding was consistent with assembly-independent, read-based profiling of phylogenetic marker genes (see **Methods**) where ∼81% of the microbial diversity was recovered in the updated MAGs (**Fig. S1B**).

To assess the taxonomic and ecological value of this new genomic resource, which we named the “Mouse B6 Gut Catalog” (MB6GC), we compared MB6GC against other murine-relevant microbiome catalogs. First, at a finer strain-level taxonomic resolution, MB6GC includes 480 medium- and high-quality MAGs (>50% completeness and <10% contamination) after de-replication at 99% ANI. Comparing these MAGs against a similar strain-resolved MAG database for CBA/J mice^29^ showed minimal overlap, with over 400 MAGs specific to C57BL/6 mice (**Fig. 1E**; see **Methods**). Second, we compared species-level (95%ANI) MAGs in MB6GC to several large murine gut metagenomic catalogs^30–32^. Despite these other catalogs integrating 3-8x more metagenomic data from 36 to 74 projects across multiple mouse strains, MB6GC added ∼6-10% new MAGs, or improved the quality (i.e., completeness and contamination) of MAGs found in the other catalogs (**Fig. 1F**; see **Methods**). Collectively, across all catalogs, 53 MAGs from MB6GC were either novel (n=15) or were the best representative genomes of their species (n=38) (**Fig. 1G**). Overall, MB6GC is a new resource to support microbiome studies in the C57BL/6 mouse strain, which is the most widely used in medical research^33^. Also, where strain-level (>99% ANI) taxonomic resolution is needed to understand the (patho)physiological significance of microbiome-host interactions^21,34^, mouse strain-specific MAG resources should be used.

### Disrupting the spinal cord-gut axis causes persistent gut dysbiosis

We next asked – ‘*Does the spinal cord regulate gut microbiome composition?*’. After data curation (details under “*Data curation*” in **Methods** and **Fig. 1A**, **Fig. S1C-D**), abundance-based beta-diversity comparisons (principal-coordinate analysis [PCoA] of Bray-Curtis dissimilarities) were made using the MAG data. These comparisons revealed sex-specific microbial communities in mice before surgery (**Fig. 2A**, **Fig. S2A**). Since sex-based differences persisted after surgery, regardless of the experimental group (**Fig. 2B**, **Fig. S2B**), we analyzed the microbiome of male and female cohorts separately. Notably, in both male and female mice, partial (T10 lesion) or complete (T4 lesion) disruption of the spinal cord-gut axis significantly changed gut microbiome composition (**Fig. 2C-D**) relative to mice with intact spinal cords (**Fig. 2E-F)**. Importantly, these changes appear to be permanent, with marked shifts in community composition (beta-diversity) evident for at least 6 months in all mice with spinal cord-gut axis dysfunction (**Fig. 3A**). Peak changes in gut microbial community composition varied by sex, occurring either three or five weeks after a partial or complete break of the spinal cord-gut axis in male or female mice, respectively (**Fig. 3A**). Like beta-diversity, disrupting spinal cord control over the gut increased alpha-diversity (estimated by Shannon’s index) in both sexes (**Fig. 3B**, **Fig. S2C**) relative to mice with intact spinal cords. An increase in alpha diversity is associated with increased gut transit time^8^, a common comorbidity of spinal cord dysfunction^35^. We confirmed this association in an independent mouse cohort using a red carmine dye assay^36^. The data show that compared to mice with intact spinal cords, total intestinal transit time increased 17-38% for at least 3 weeks in mice with a spinal lesion (**Fig. 3C**).

**Fig. 2.**
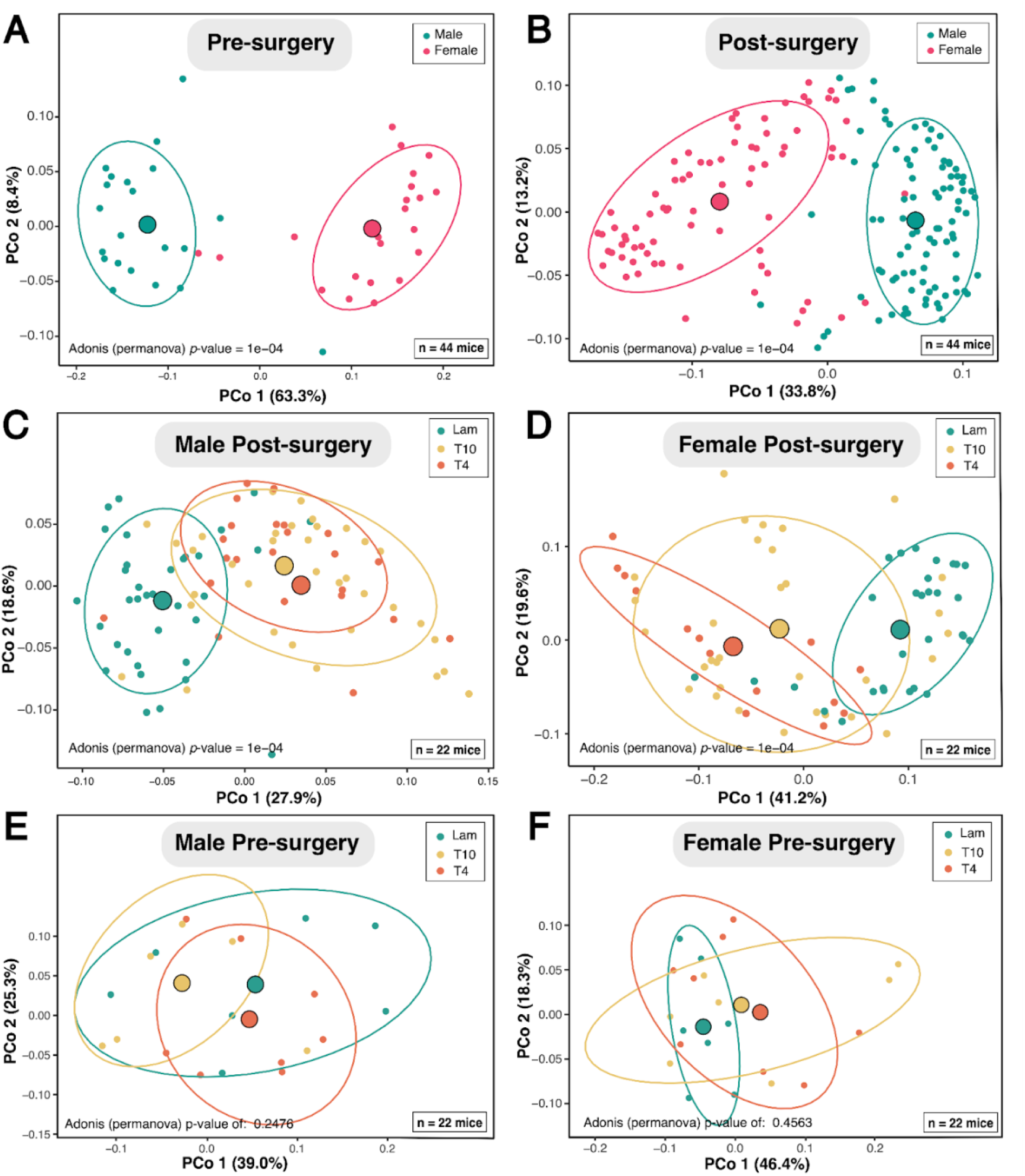
Community-level microbiome changes are spinal cord-dependent and vary as a function of sex and spinal lesion level. (**A-B**) MAG abundance-based principal coordinate analysis (PCoA) of Bray-Curtis dissimilarities from metagenomes of 44 mice (see “*Data Curation”* in **Methods** and **Fig. S1C**) show sex-based differences before (**A**) and after (**B**) spinal surgery. Consistent results arise from analyses before data curation (i.e., including outliers; see **Fig. S2A-B**). (**C-F**) PCoAs constructed using the same methods as in (**A-B**) show microbiome differences only after a break in the spinal cord-gut axis (**C-D**). Before spinal lesioning (**E-F**) no group differences exist in male (n=22) or female (n=22) mice. See permanova *p*-values for assessment of significant differences in each panel. Small dots represent individual samples, while large dots represent the centroids per sex (**A-B**) or surgery (**C-F**).

**Fig. 3.**
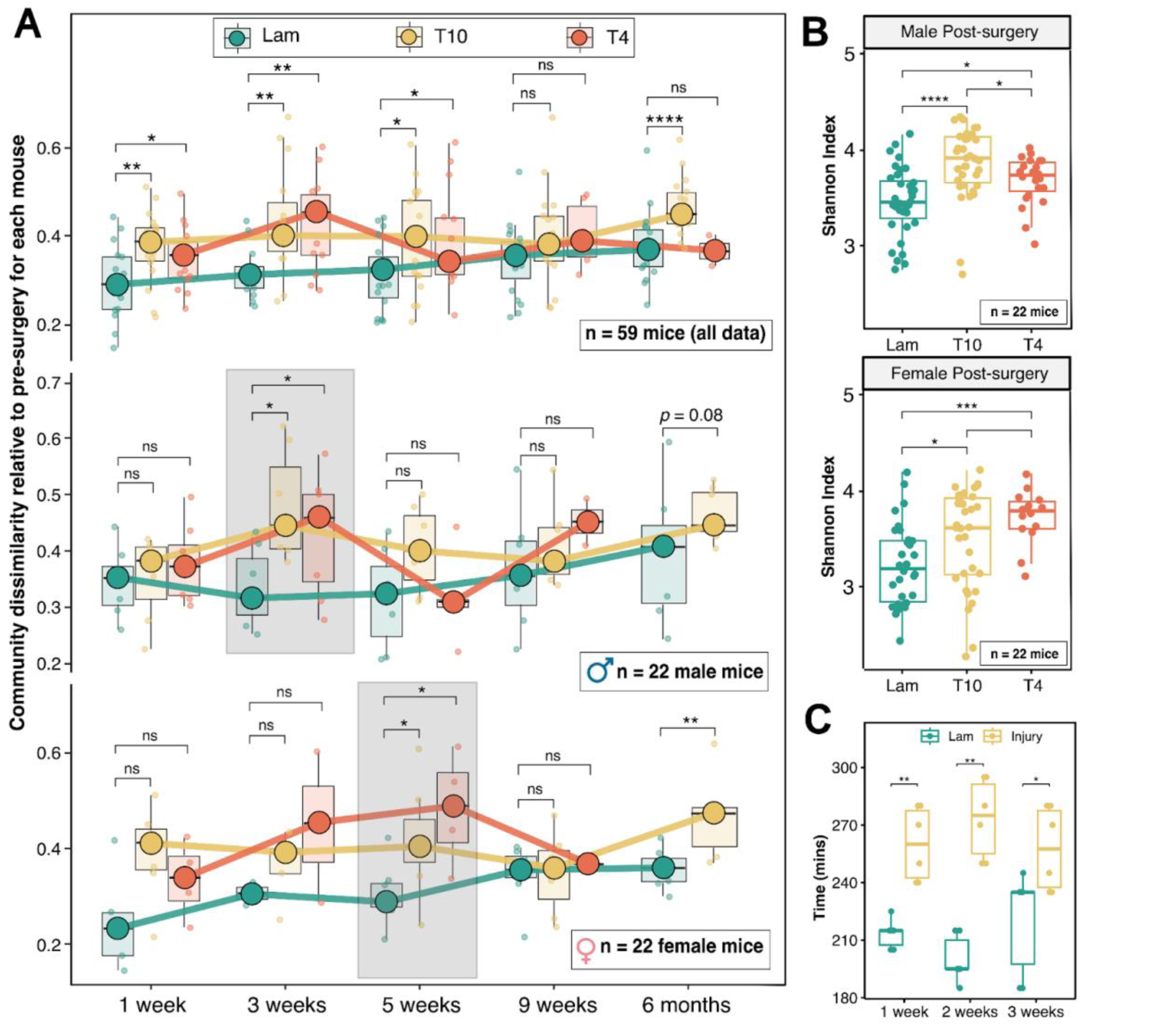
Dynamic changes in microbiome beta- and alpha-diversity are spinal cord-dependent and vary as a function of sex and spinal lesion level. **(A)** Boxplot injury-level- and time-resolved comparisons of Bray-Curtis dissimilarities (beta-diversity) relative to pre-surgery microbiome, shown for all data (n=59 mice) as well as curated male (n=22) and female (n=22) mouse data (see “*Data curation”* in **Methods** and **Fig. S1C**). Each small dot represents the microbiome change of an independent mouse relative to its baseline microbiome. Boxplot median values across mice per time-point are represented by large dots and are connected with thick lines. Peak changes are highlighted with gray boxes in male (3 weeks) and female (5 weeks) mice. (**B**) Boxplots showing spinal injury-level dependent microbiome alpha-diversity changes in curated male (n=22) and female (n=22) mice data. See baseline (Day 0) and sex- and time-resolved alpha diversity patterns in **Fig. S2C**. (**C**) Complete spinal injury slows intestinal transit time (Red carmine dye assay) relative to sham-lesion (Lam) controls. For all boxplots, whiskers represent the 25th and 75th percentile ranges while *p*-values represent those from Wilcoxon tests, adjusted for multiple comparisons by the fdr (false discovery rate) method (see **Methods**). ****: *p* <= 0.0001, ***: *p* <= 0.001, **: *p* <= 0.01, *: *p* <= 0.05, ns: *p* > 0.05.

Overall, instead of static, snapshot comparisons of microbiomes from different mice after surgery, which is the norm in most microbiome studies, here, within-mouse longitudinal comparisons (i.e., before and periodically after injury) (**Fig. 3**) reveal that spinal cord dysfunction causes persistent and dynamic changes in gut microbiome composition. Moreover, our study confirms that sex is a critical biological variable when analyzing the microbiome (see also^37,38^). Thus, an intact spinal cord is required for regulating gut microbiome composition, regardless of sex. Notably, prior animal studies have mostly interrogated the effects of spinal cord dysfunction on gut microbiome composition in female mice only (reviewed in^16^). Sex-specific differences in the microbiome are linked to development of, or susceptibility to, autoimmune disease^39–41^, gastrointestinal disease^42,43^, metabolic disease^44,45^, neuropsychiatric and neurodegenerative disease^46,47^. Collectively, our analyses extend existing microbiome datasets to include longitudinal spinal cord-dependent control over the gut ecosystem in both sexes, providing both a valuable new experimental resource and a more comprehensive understanding of CNS regulation of the microbiome.

### Microbes that persist after disrupting the spinal cord-gut axis include taxa previously linked to the host physiological state

Next, to determine which taxa respond to spinal cord-gut axis perturbation, microbial relative abundances were compared in mice with or without an intact spinal cord (see **Methods** for details on statistical analysis). Significant spinal cord-dependent effects are shown at an aggregated phylum-level (**Fig. 4A**, **Fig. S3A**), and an individual MAG/species-level (**Fig. 4B-C**, **Fig. S3B**). At the phylum level, a break in the spinal cord-gut axis decreased Bacteroidota and Proteobacteria, with a concurrent increase in Firmicutes_A in males and females (**Fig. 4A, Fig. S3A**). Whereas the ratio between the phyla Firmicutes and Bacteroidetes was also reported to increase in our earlier work in mice with spinal cord injury^24,48^, others have reported that this same ratio decreases^49,50^. This discrepancy may be attributed to improvements in or differences between databases, as the GTDB uses genomic data that now distinguish the Firmicutes phyla into three separate phyla: Firmicutes, Firmicutes_A, and Firmicutes_B (see “*GTDB vs NCBI Taxonomy*” in **Methods**). In addition, other lower-resolution techniques (i.e., 16S rRNA gene amplicon sequencing) may not be able to distinguish between these three classically lumped lineages. Here, using genome-resolved metagenomics, Firmicutes_A was the only phylum that changed among the three.

**Fig. 4.**
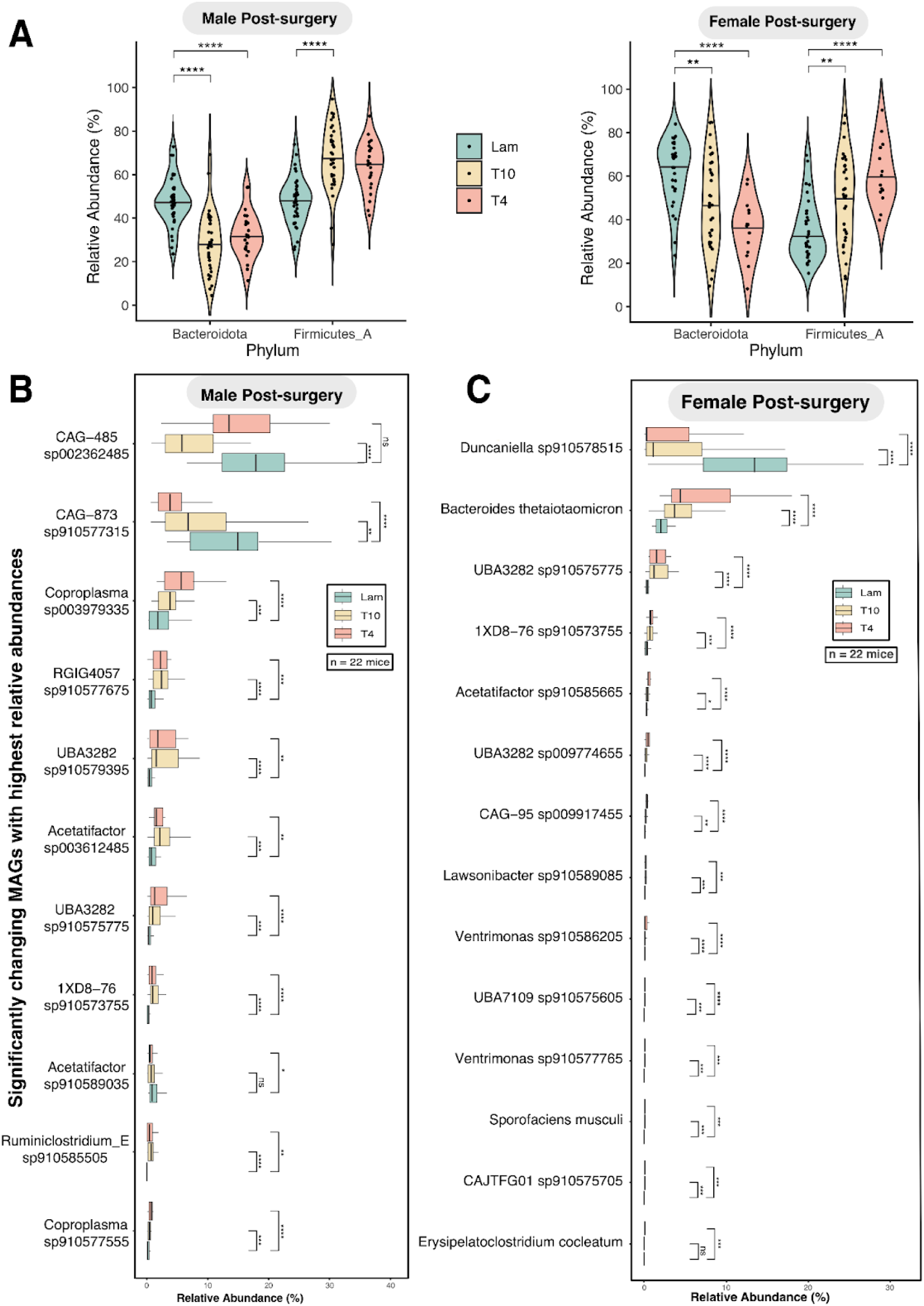
Significant phylum- and MAG-level microbiome changes occur after disrupting the spinal cord-gut axis. **(A)** Violin plots of the most abundant (median relative abundance >0.25%) phyla show that Bacteriodota and Firmicutes_A significantly change only after disrupting the spinal cord-gut axis in male (right) and female (left) mice. Pre-surgery control plots and plots of the least abundant phyla (i.e., <0.25 median relative abundance) can be found in **Fig. S3A**. (**B**) Boxplots of the most abundant (median relative abundance > 0.5%) significantly changing MAGs in male mice with spinal lesions. All 68 changing MAGs can be found in **Fig. S3B**. (**C**) Boxplots of all significantly changing MAGs (n=14) in female mice with spinal lesions. For violin plots and boxplots in all panels (**A-C**), the 25th and 75th percentile ranges are shown. *p*-values represent those from Wilcoxon tests, adjusted for multiple comparisons by the fdr (false discovery rate) method (see “*Statistical methods for determining differentially abundant taxa*” in **Methods**). ****: *p* <= 0.0001, ***: *p* <= 0.001, **: *p* <= 0.01, *: *p* <= 0.05, ns: *p* > 0.05.

At the species level, many differentially abundant MAGs were found in male (n=68; **Fig. 4B, Fig. S3B**) and female (n=14; **Fig. 4C**) mice. The magnitude of change in relative abundance of several discrete microbial species (**Fig. 4B-C**, **Fig S3B**) was differentially affected by spinal lesion level, an observation that was also shown for species alpha diversity (**Fig. 3B**). Temporal changes in MAG abundances also varied by spinal lesion level and sex (**Fig. S4**). However, time-resolved analysis of rare taxa shows persistent spinal lesion-dependent changes in *Lactobacillus johnsonii* (**Fig. 5A**) in both sexes. Notably, changes in *Weissella cibaria* (a significantly changing taxon in^24^) were time-dependent (**Fig. S4G**), highlighting the importance of well-controlled temporal analyses to discern changes in the gut microbiome that may only be sporadically evident. Indeed, static, time-delimited sampling at any single time point would likely under- or overestimate change(s) for a given microbe or the effect that sex or the spinal cord have on these changes.

**Fig. 5.**
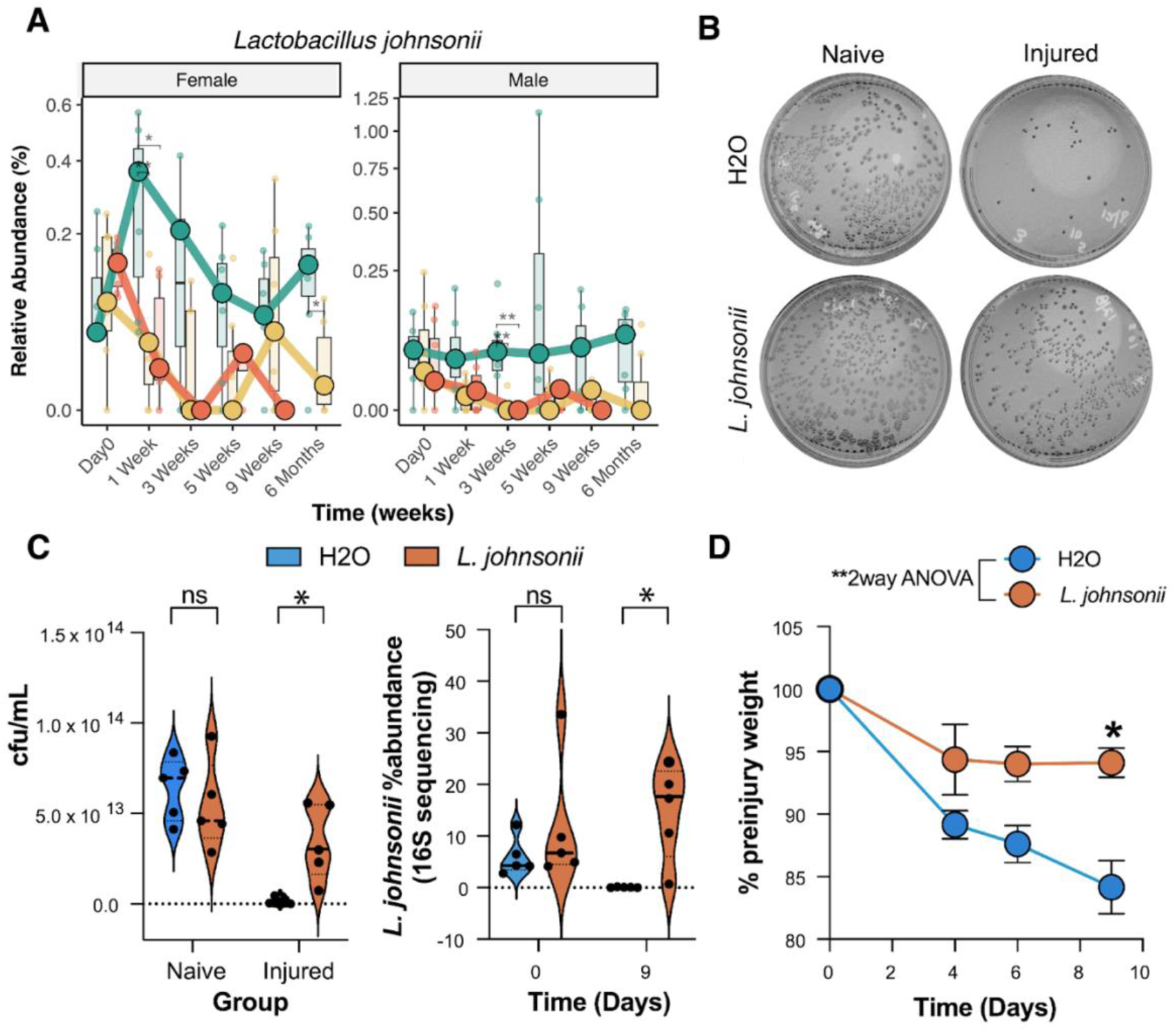
The effects of feeding *L. johnsonii* to overcome the effects of a break in the spinal cord-gut axis. **(A)** A rapid and protracted reduction in *L. johnsonii* MAG abundance occurs in the gut of male and female mice after a break in the spinal cord-gut axis. (**B-C**) Fecal cultures grown on MRS agar show reduced *L. johnsonii* colonies in mice subjected to a clinically-relevant model of spinal cord injury (SCI). SCI-dependent loss of *L. johnsonii* can be reversed by feeding mice 1 x 10^8^cfu *L. johnsonii* daily after SCI **(C left**). 16S sequencing of fecal samples confirmed that this feeding regimen increased the abundance of *L. johnsonii* in the gut (**C right**). (**D**) Treatment with an *L. johnsonii* “probiotic” attenuates SCI-induced weight loss. *p*-values in (**A**) represent those from Wilcoxon tests, adjusted for multiple comparisons by the fdr (false discovery rate) method (see “*Statistical methods for determining differentially abundant taxa*” in **Methods**). ****: *p* <= 0.0001, ***: *p* <= 0.001, **: *p* <= 0.01, *: *p* <= 0.05, ns: *p* > 0.05. *p*-values for (**C-D**) are from 2-way ANOVA with Šídák’s multiple comparisons test (**C**) or repeated measures 2-way ANOVA with Bonferroni’s multiple comparisons test (**D**).

Although abundant MAGs that decrease post-lesion in males and females were mostly Muribaculaceae family members (3 out of 4), most increasing MAGs (14 of 21) belong to the Lachnospiraceae family (**Table S1**). A decrease in Muribaculaceae abundance with a concomitant increase in Lachnospiraceae has been linked to metabolic diseases, such as obesity, type 2 diabetes, insulin resistance, and non-alcoholic steatohepatitis^51,52^. Other rare taxa that are reduced in mice without normal spinal cord control of the gut, including *L. johnsonii* and Turicibacter sp002311155, may also play critical roles in influencing mammalian physiology and health^53–55^. Because *L. johnsonii*’s MAG was essentially depleted after lesioning the spinal cord, regardless of sex or spinal lesion level (**Fig. 5A**), and because this microbe was previously identified as a potential beneficial microbe capable of regulating various murine metabolic pathways^24^, we directly tested the effects of reintroducing *L. johnsonii* into the gut of mice with spinal cord lesions.

### Spinal cord injury-dependent weight loss is prevented by post-injury feeding with *Lactobacillus johnsonii*

We isolated *L. johnsonii* by culturing fecal samples from naive C57BL/6 mice on a selective growth medium for *Lactobacilli* (see **Methods**). Female C57BL/6 mice received complete mid-thoracic spinal lesions (injured) or were left uninjured (naive). Starting at 1 day post-lesion, naive and injured mice were fed 1 x 10^8^ cfu of *L. johnsonii* or vehicle (water) daily via oral gavage. At 9 days post-lesion, fecal samples from all mice were plated then cfu/ml was calculated. A significant decrease in *L. johnsonii* was noted in injured mice fed water only (**Fig. 5B**), confirming metagenomic data from this and our prior study (**Fig. 5A**; and^24^). Conversely, feeding *L. johnsonii* significantly increased its abundance in lesioned mice, which was visually evident in cultures, confirmed by quantifying cfu/ml and by 16S sequencing of fecal samples obtained from mice in each cohort (**Fig. 5B-C**). Importantly, rapid weight loss, a consistent phenomenon indicative of a changing metabolic state in mice with spinal injuries, was significantly reduced when mice were fed *L. johnsonii* (**Fig. 5D**). Since feeding *L. johnsonii* did not affect weight in naive mice (not shown), we conclude that these results reflect a gut-microbiome-dependent “rescue” of metabolic perturbations caused by a break in the spinal cord-gut axis. For example, spinal cord injury causes rapid muscle wasting^56^, and *Lactobacillus* can prevent sarcopenia by modulating inflammation, oxidative stress, and skeletal muscle metabolism^57^. Although further research is needed to determine the precise mechanisms underlying the rescue effects of *L. johnsonii*, this bacterium can improve intestinal barrier function, modulate systemic immunity, and prevent metabolic disease^53,58–61^.

### Altered carbohydrate metabolism in spinal cord-dependent dysbiotic gut selects against rare beneficial taxa (e.g. *L. johnsonii*)

Data in **Fig. 5** demonstrate the physiologic importance of *L. johnsonii*, especially when the spinal cord-gut axis is perturbed, as is common in most types of neurological disease. But, why would a sustained reduction in *L. johnsonii* occur in these disease states? To answer this question, we used the entire MB6GC to infer the complex metabolic interactions that occur across the gut microbial community, focusing on the MAGs (n=13) that were most abundant and changing after a break in the spinal cord-gut axis (**Fig. 4B-C**).

All MB6GC MAGs were annotated using DRAM (Distilled and Refined Annotation of Metabolism^62^), a tool designed to functionally annotate microbial genes and categorize them across various expert-curated metabolic pathways. Interestingly, the four most abundant MAGs in non-lesioned mice (collectively ∼28% and ∼45% median relative abundance in female and male mice, respectively) belong to the Bacteroidales order (**Table S1**), namely CAG-485 sp002362485, CAG-873 sp910577315, Duncaniella sp910578515, and *B. thetaiotaomicron*. Their proportionally large relative abundance can be attributed to these bacteria encoding a complete electron transport chain, including the high-oxygen affinity cytochrome *bd* (**Fig. S5**), allowing them to survive and potentially thrive in the low oxygen environment of the gut^63^. Additionally, under homeostatic conditions, species from the Bacteroidales order can perform polysaccharide to oligo-and monosaccharide conversions that dictate cross-feeding relationships broadly across a gut microbial community^34,51,64^. The products of these carbohydrate conversions are primary energy sources for many gut bacteria, and carbohydrate metabolism by gut microbes is essential for producing various beneficial metabolites (e.g. short-chain fatty acids; SCFAs) that support host health^34^. However, as mechanistic genome-resolved approaches are only just emerging in studies of mammalian gut microbiome, community-contextualized microbial carbohydrate cross-feeding after dysbiosis has not been well-resolved in any system, especially involving *L. johnsonii*. Here, we examine state-dependent changes in systems used by bacteria to degrade and take up carbohydrates, namely glycoside hydrolases (GHs), sugar transporters, and polysaccharide utilization loci (PUL) systems.

Based on the 13 most abundant changing MAGs, the capacity to degrade complex polysaccharides remained unchanged after disrupting the spinal cord-gut axis. This conclusion was drawn from analyzing the number and types of GHs, enzymes responsible for breaking down complex carbohydrates into simple sugars for microbial energy. When GHs were grouped into 14 substrate-based categories (*sensu*^62^), 11 of 13 MAGs encoded diverse GHs that act on all 14 carbohydrate substrates (**Fig. S6A**). Of these 11 MAGs, three decreased and eight increased in mice with spinal cord lesions (**Fig. S6B**). Unlike these abundant taxa, which can degrade these 14 carbohydrates, *L. johnsonii* scarcely encodes GHs (**Fig. S6A**; see also *Turicibacter* sp002311155, another rare beneficial taxon that is decreased in mice with spinal lesions^53–55^). These data suggest that rare beneficial taxa depend on abundant community members for polysaccharide degradation and cross-feeding. Indeed, MAGs of these rare taxa encode many diverse types of sugar transporters that would enable them to scavenge simple sugars (**Fig. S7**). This suggests that, under homeostasis, the final products of polysaccharide degradation carried out by abundant community members (e.g., Muribaculaceae) might be used by rare, beneficial taxa. After a significant perturbation, as Muribaculaceae decrease, the increase in other abundant bacteria is unlikely to support cross-feeding of rare taxa, a potential mechanism causing prolonged dysbiosis. For example, Lachnospiraceae family members encode GHs with broad polysaccharide substrate specificity (**Fig. S6**), but unlike Muribaculaceae, Lachnospiraceae also encode other numerous and diverse simple sugar transporters (**Fig. S7**), so they are expected to consume the simple sugar products of polysaccharide degradation. Conversely, the four most abundant MAGs changing post-lesion (all Bacteroidales) scarcely encode these sugar transporters (**Fig. S7**), and instead exclusively encode polysaccharide utilization loci (PULs), with multiple characteristic homologs of the starch utilization system (**Fig. S8**). Bacteroidales use these systems to sense, bind, degrade, and import polysaccharides and oligosaccharides^65^. Of these four Bacteroidales MAGs, one increases after spinal lesioning (*B. thetaiotaomicron*) and three less-characterized Muribaculaceae decrease (**Fig. 4 and Fig. S4**), with no evidence for such systems in any other taxa, consistent with previous reports^65^.

PUL systems exhibit selfish and altruistic behaviors in complex trophic networks like in the gut. Some Bacteriodales species encode “selfish” PUL systems that rapidly import broken-down oligosaccharides into the periplasm for further breakdown, conferring no direct benefit to neighboring species, while other Bacteroidales encode “leaky” PULs that release partial breakdown products (PBPs) for the benefit of the whole community^66^. We found evidence for a selfish PUL system (utilizing GH99 and GH76; reviewed in^66^) encoded only by *B. thetaiotaomicron*, potentially explaining the relative fitness of this taxon in the dysbiotic gut of mice with spinal lesions, where transporter-rich Lachnospiraceae compete for and scavenge monosaccharides. Other Bacteroidales species, notably those of the Muribaculaceae family, exhibit an “altruistic” behavior by selectively packaging GHs into outer membrane vesicles that are released into the extracellular environment, enabling the production of free saccharides for the greater gut community^66–68^. More broadly, the Muribaculaceae family has been credited with probiotic roles, including a protective oxygen-scavenging role^51^ – consistent with the cytochrome *bd* results above - see **Fig. S5**) and cross-feeding with other microbial species (reviewed in^51^). Indeed, we find that some Muribaculaceae taxa (e.g., CAG-485 sp002362485) encode diverse GHs, potentially boosting the fitness of other Bacteroidales (e.g, CAG-873 sp910577315) and rare beneficial taxa (e.g., *L. jonhsonii*) that encode scarce GHs and instead depend on PBPs or monosaccharide products, respectively.

A conceptual model that explains gut ecosystem function, both before and after a break in the spinal cord-gut axis, can be developed from our data (**Fig. 6**). Fundamental to this model is our data-driven hypothesis that reassembly of the microbial community post-injury, as orchestrated by the most abundant and dynamic taxa, results in a unique carbohydrate metabolic fingerprint that disfavors rare beneficial taxa and cooperative trophic interactions. First, under homeostatic conditions, some abundant Bacteroidales members break down polysaccharides into oligo- and mono-saccharides, “public goods” that other bacteroidales and beneficial microbes, such as *L. johnsonii* and *Turicibacter* sp002311155, take up for energy using their sugar transporters. Second, after a break in the spinal cord-gut axis, the niche of Bacteroidales (Muribaculaceae members and *B. thetaiotaomicron*) is partially filled by other polysaccharide degraders, notably Lachnospiraceae family members. While Muribaculaceae MAGs significantly decrease, *B. thetaiotaomicron* seem to thrive, possibly because their PUL systems do not discard resources for other members and instead can quickly import oligosaccharides back to the periplasm. Unlike Bacteroidales, other polysaccharide degraders, mostly Lachnospiraceae family members, also encode large numbers of diverse sugar transporters and thus can fully utilize carbohydrates and consume the monosaccharides they produce, essentially starving rare beneficial taxa (e.g., *L. johnsonii*) and reducing their abundance in the community (**Fig. 6**).

**Fig. 6.**
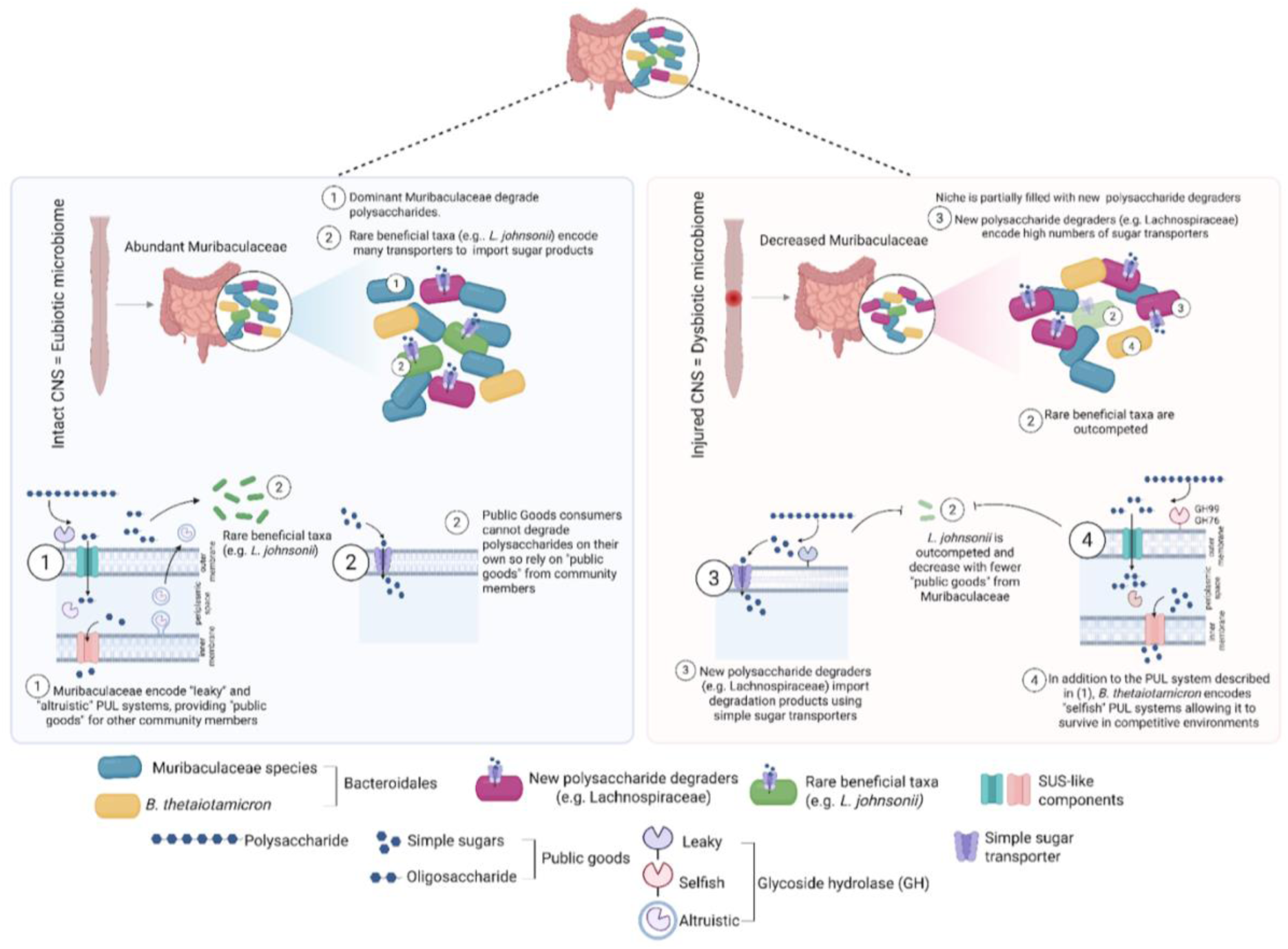
Data-driven model of gut microbial community carbohydrate metabolism explains *L. johnsonii* depletion in mice with spinal lesions. Metabolic cross-feeding model shows that a decrease in Muribaculaceae likely results in fewer public goods available to rare beneficial taxa like *L. johnsonii*. As Muribaculaceae decrease, their niche is partially replaced by other polysaccharide degraders that import most of the oligo- and mono-saccharides for their own use (e.g. Lachnospiraceae). This effectively starves rare beneficial taxa that cannot compete with taxa that encode “selfish” carbohydrate degradation and import systems.

This model of gut ecosystem function provides a framework for predicting changes in gut microbe-host interactions beyond disruptions in the spinal cord-gut axis. For example, a study using paired metagenomics and metabolomics in insulin-resistant individuals (characterized by elevated sugar levels in blood, urine, and feces) found that Lachnospiraceae taxa positively correlate with monosaccharide concentrations in fecal matter, whereas Bacteroidales taxa show negative correlations with the same metabolites^69^. Our model suggests that high monosaccharide concentrations favor transporter-rich taxa like Lachnospiraceae while disfavoring transporter-scarce taxa like Bacteroidales.

## Conclusions

This report establishes the spinal cord as an integral but largely overlooked part of the “CNS-gut-microbiota axis”. By enhancing study design, sample diversity, and analytical methods, we introduce new resources (e.g., MB6GC) and microbial targets for better understanding disease mechanisms and potential interventions. For instance, feeding *L. johnsonii* helps restore host metabolic benefits during recovery from spinal cord injury (**Fig. 5**). A grand challenge in microbiome science is deciphering ecosystem-relevant changes in microbial functions without prior knowledge bias. Applying ecological theory and eco-systems biology concepts can organize significant taxa (∼70 MAGs here) into key sub-communities controlled by a few dominant taxa, which represent priority targets for mechanistic studies. To achieve this and further reveal the scope and potential biological consequences of a break in the CNS-gut-microbiota-gut axis, a time-resolved, ecosystem-aware workflow is necessary to link host and gut microbiome metabolisms.

**Fig. S1.**
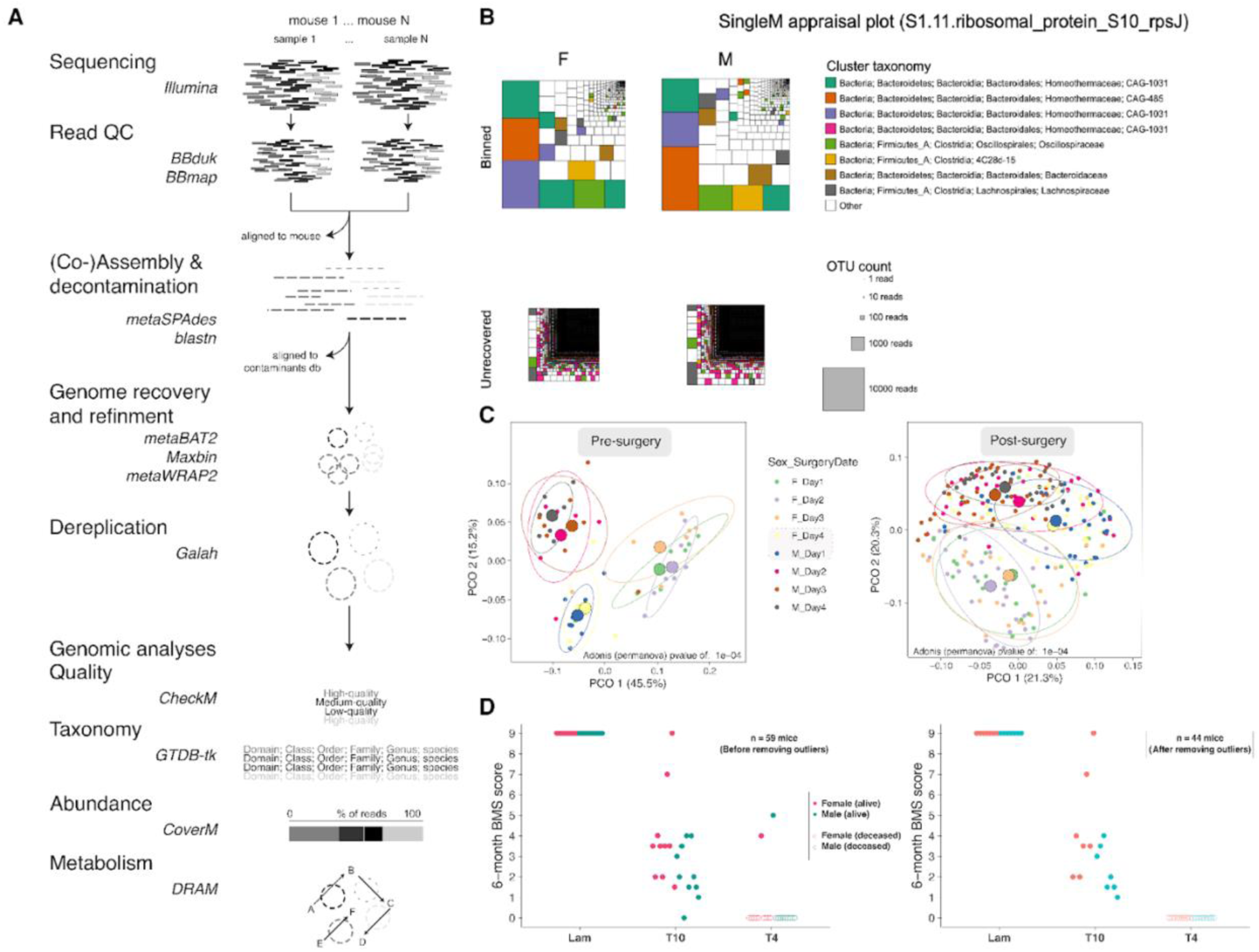
Overview of the bioinformatic pipeline, read-profiling, and data curation. **(A)** Overview of the bioinformatic pipeline used in this study for metagenomic analysis of gut microbiomes. Following sequencing, read quality control (QC) was carried out including read trimming and removal of reads matching to the mouse genome. Co-assembly of QC’d reads per mouse was done, followed by decontamination of scaffolds to be used for binning. Genome recovery, refinement, and dereplication at 95% ANI resulted in 263 MAGs that were used for further analysis. (**B**) read-based profiling of phylogenetic marker genes (see “*Read-based profiling*” in **Methods**) from raw reads suggest that, in both male and female mice, ∼81% of the microbial diversity in the ecosystem was recovered in our updated MAG database. (**C**) Principal coordinate analysis (PCoA) based on Bray-Curtis dissimilarities for pre-surgery (left = Day 0) and post-surgery (right) microbiome samples, colored based on the combination of sex and surgery date (see “*Data curation*” in **Methods**) to discern confounders. Male mice having their operation performed on the first surgery day and female mice having theirs at the fourth day clustered closer to each other and differently from the larger male and female clusters, as indicated by a significant separation pre- and post-surgery (adonis *p*-value = 1e-04). (**D**) Basso Mouse Scale (BMS) scores taken at 6 months post-injury after data curation (n = 44 mice) – see the caption of **Fig. 1**.

**Fig. S2.**
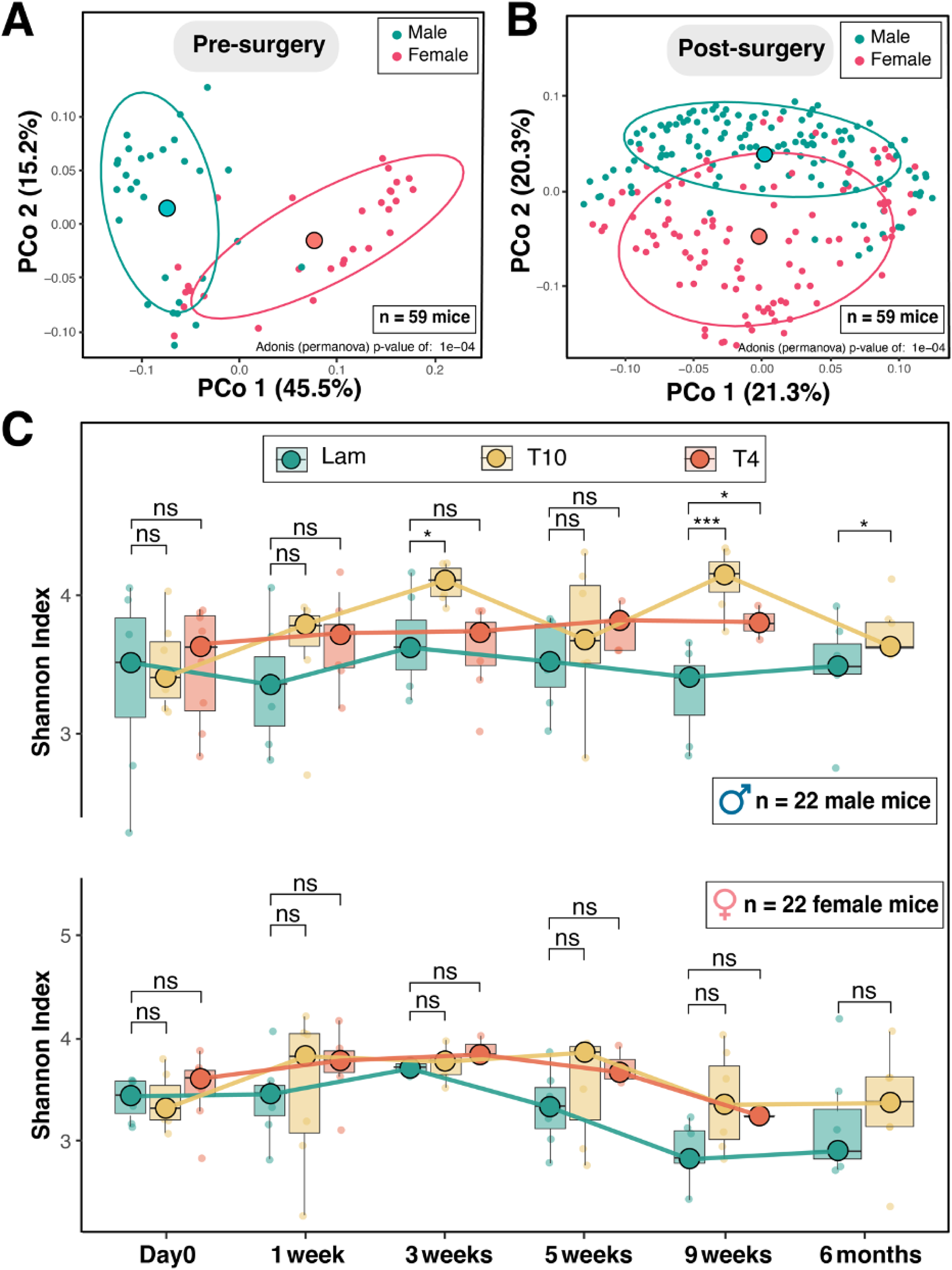
PCoAs for male and female and alpha diversity. (**A-B**) MAG abundance-based principal coordinate analysis (PCoA) of Bray-Curtis dissimilarities from metagenomes of 59 mice (including outliers), showing sex-based differences before (**A**) and after (**B**) surgery. See permanova *p*-values for assessment of significant differences in each panel. Small dots represent individual samples while large dots represent the centroids per sex. (**C**) Boxplot injury-level- and time-resolved comparisons of Shannon’s index, shown for curated male (n=22) and female (n=22) mice data. Each small dot represents the microbiome changes of an independent mouse at that timepoint. Boxplot median values across mice per timepoint are represented by large dots and are connected with thick lines. This panel provides a detailed temporal overview of the collective changes shown in **Fig. 3**, highlighting changes at specific time points and the apparent reduction in statistical power for per-timepoint analyses, especially in female mice. For all boxplots, whiskers represent the 25th and 75th percentile ranges while *p*-values represent those from Wilcoxon tests, adjusted for multiple comparisons by the fdr (false discovery rate) method (see **Methods**). ****: *p* <= 0.0001, ***: *p* <= 0.001, **: *p* <= 0.01, *: *p* <= 0.05, ns: *p* > 0.05.

**Fig. S3.**
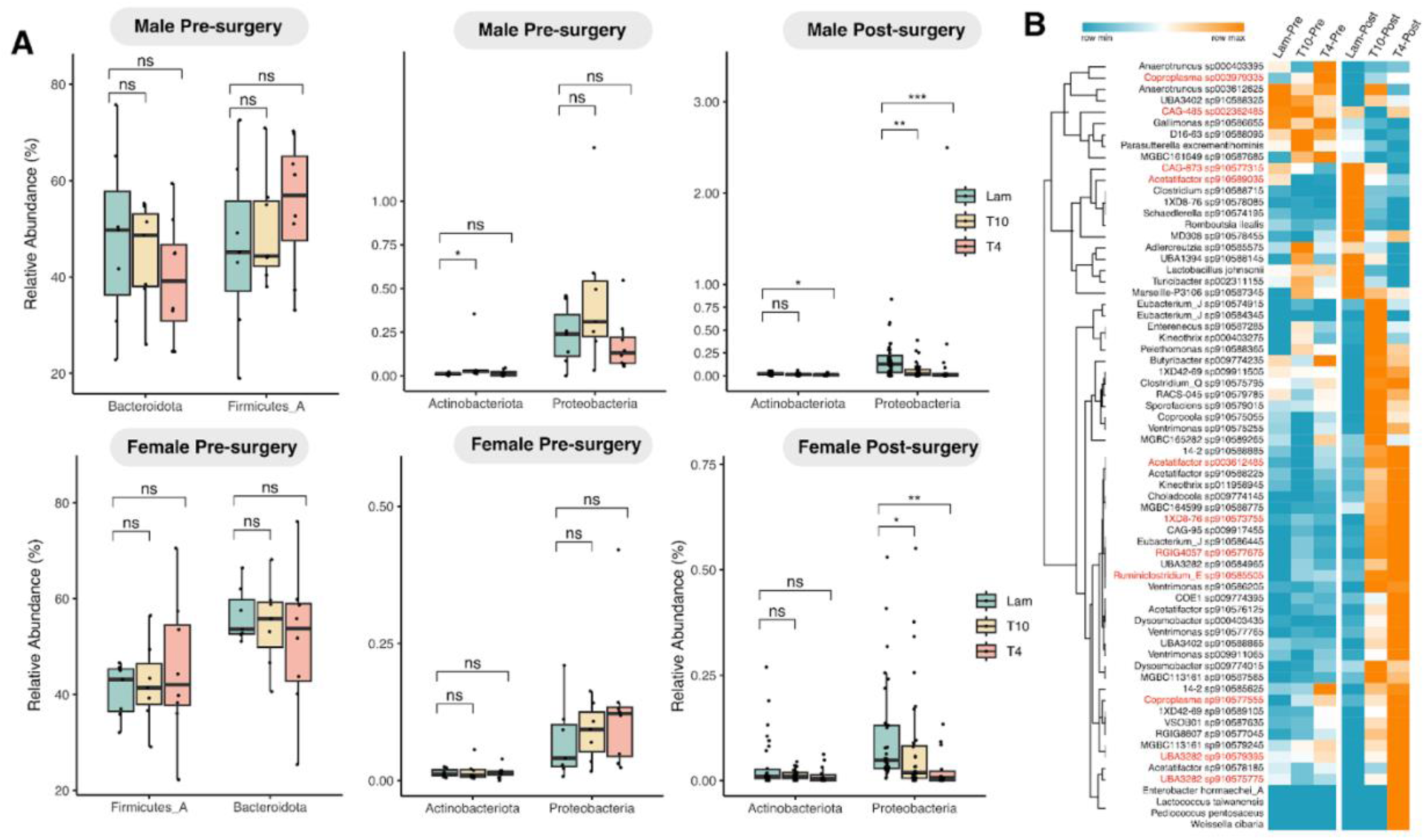
Other significant phylum- and MAG-level microbiome changes after disrupting the spinal cord-gut axis (n=22). (**A left**) Boxplots of the most abundant phyla, Bacteroidota and Firmicutes_A at baseline (Day0) for male (top) and female (bottom) mice; (**A middle**) Boxplots of the least abundant phyla, Actinobacteriota and Proteobacteria, at baseline in male (top) and female (bottom) mice; (**A right**) Boxplots of the least abundant phyla, Actinobacteriota and Proteobacteria, after surgery in male (top) and female (bottom) mice. (**B**) Heatmap of z-score-normalized MAG mean relative abundances before and after lesioning the spinal cord in male mice. Only significantly changing MAGs (n=68) are shown here. MAGs with the highest median relative abundances are highlighted in red and further shown in **Fig. 4B**. For boxplots in panel (A), the 25th and 75th percentile ranges are shown. *p*-values represent those from Wilcoxon tests, adjusted for multiple comparisons by the fdr (false discovery rate) method (see “*Statistical methods for determining differentially abundant taxa*” in **Methods**). ****: *p* <= 0.0001, ***: *p* <= 0.001, **: *p* <= 0.01, *: *p* <= 0.05, ns: *p* > 0.05.

**Fig. S4.**
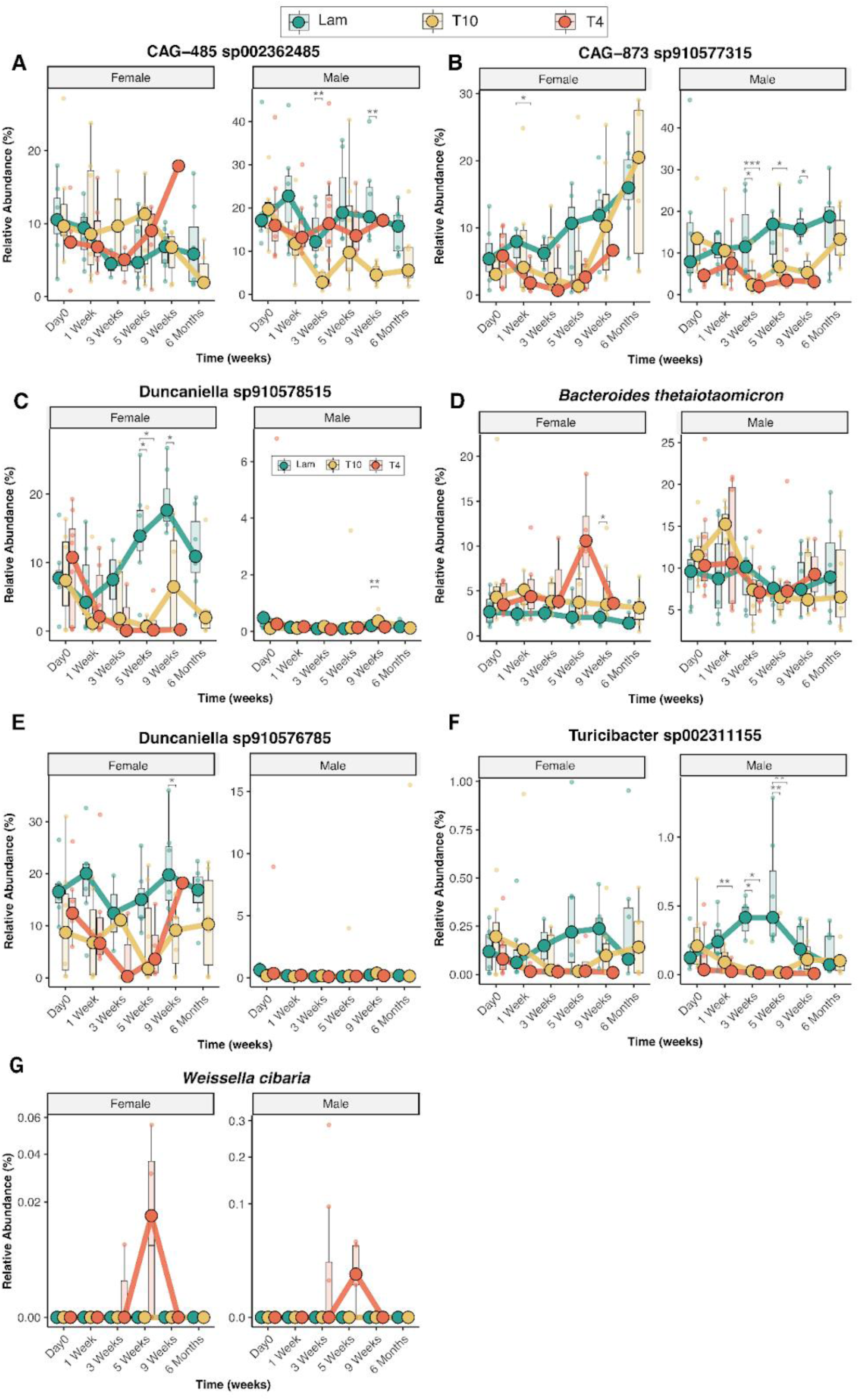
Temporal MAG abundance dynamics across the three experimental groups. Boxplots showing temporal dynamics of abundant bacteroidales MAGs (**A-E**), the MAG for the rare beneficial Turicibacter taxon (**F**), and the sporadically changing *Weisella cibari* MAG (**G**) in female (left insets) and male (right insets) mice resolved by experimental group and individual samples. The most abundant differentially affected MAGs in male (i.e., CAG-485 sp002362485 and CAG-873 sp910577315 shown in **A-B**) and female (Duncaniella sp910578515 shown in **C**) mice belong to the Muribaculaceae family, with persistent, spinal cord-dependent reductions in Duncaniella sp910578515 noted in female, but not male mice, and mixed effects on CAG-485 sp002362485. Spinal-dependent changes in *Bacteroides thetaiotaomicron* (**D**) occurred in female mice only. Some differentially abundant species identified post-lesion were reported in our earlier pilot study^24^, such as CAG-1031 bins – here CAG-1031 bins have been reclassified to Duncaniella sp910578515 and Duncaniella sp910576785 (**D-E**) – and other rare taxa including *Lactobacillus johnsonii* (**Fig. 5A**), *Weissella cibaria* (**F**), and the Turicibacter genus (its MAG shown in **G**). Each small dot represents a MAG relative abundance at a specific time-point from a single mouse over 6 months. Boxplot median values across mice per time-point are represented by large dots and are connected with thick lines. For all boxplots, whiskers represent the 25th and 75th percentile ranges while *p*-values represent those from Wilcoxon tests, adjusted for multiple comparisons by the fdr (false discovery rate) method (see **Methods**). ****: *p* <= 0.0001, ***: *p* <= 0.001, **: *p* <= 0.01, *: *p* <= 0.05, ns: *p* > 0.05.

**Fig. S5.**
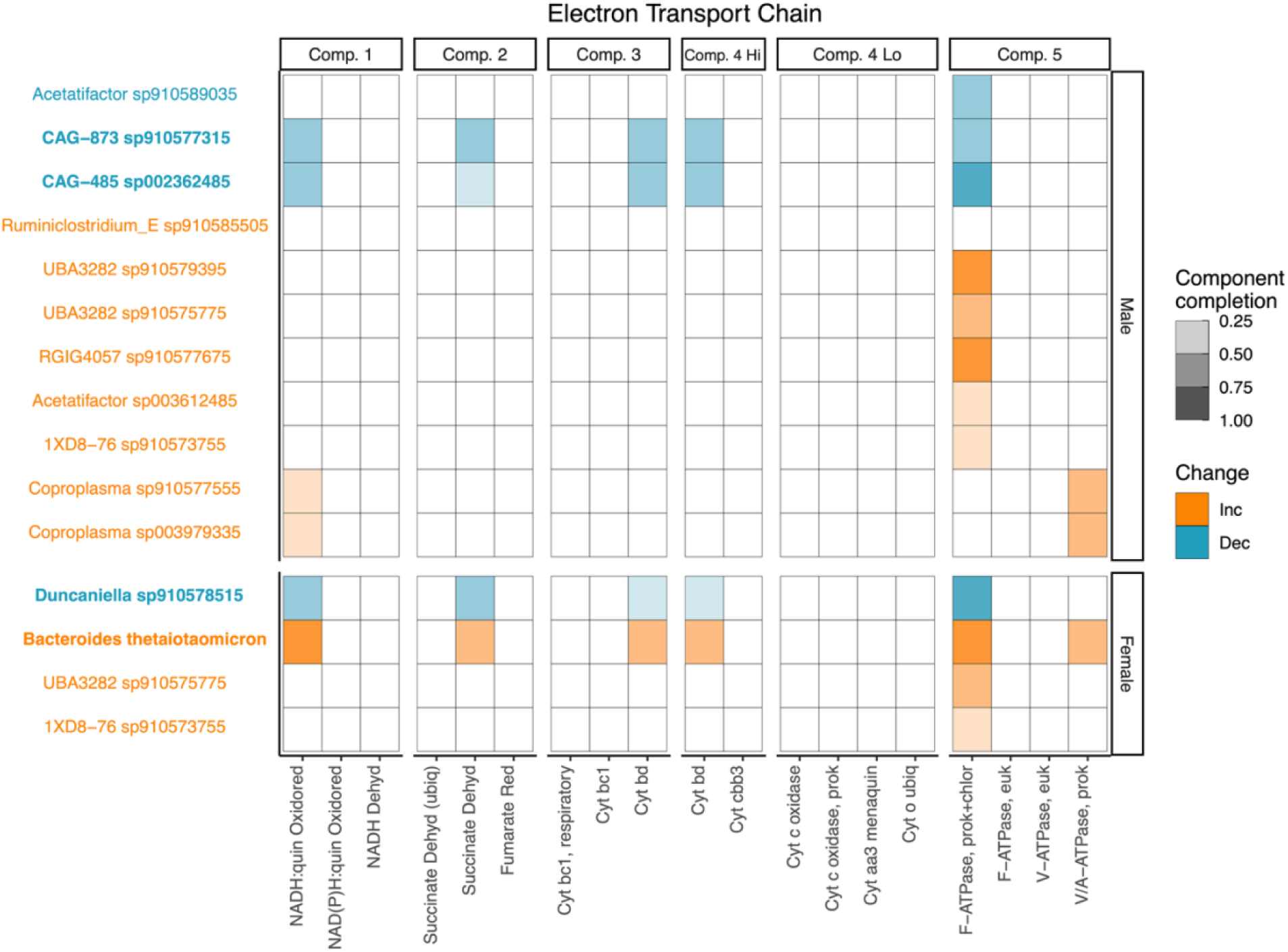
Electron transport chain (ETC) components encoded by the most abundant changing MAGs in male and female mice. A heatmap showing the completeness of each component of the ETC encoded by the most abundant changing MAGs according to annotations by DRAM. The color indicates the direction of change (orange for increasing and blue for decreasing) after a break in the spinal cord-gut axis while color intensity indicates component completion. MAGs in bold encode the minimum number of components required for a complete ETC. For example, a MAG encoding cytochrome *bd* or cytochrome *cbb3* is capable of respiring oxygen via their high-affinity cytochrome (Comp. 4 Hi). None of the MAGs encoded the low oxygen-affinity cytochromes (Comp. 4 Lo) while bacteroidales encoded the high oxygen-affinity cytochrome *bd*.

**Fig. S6.**
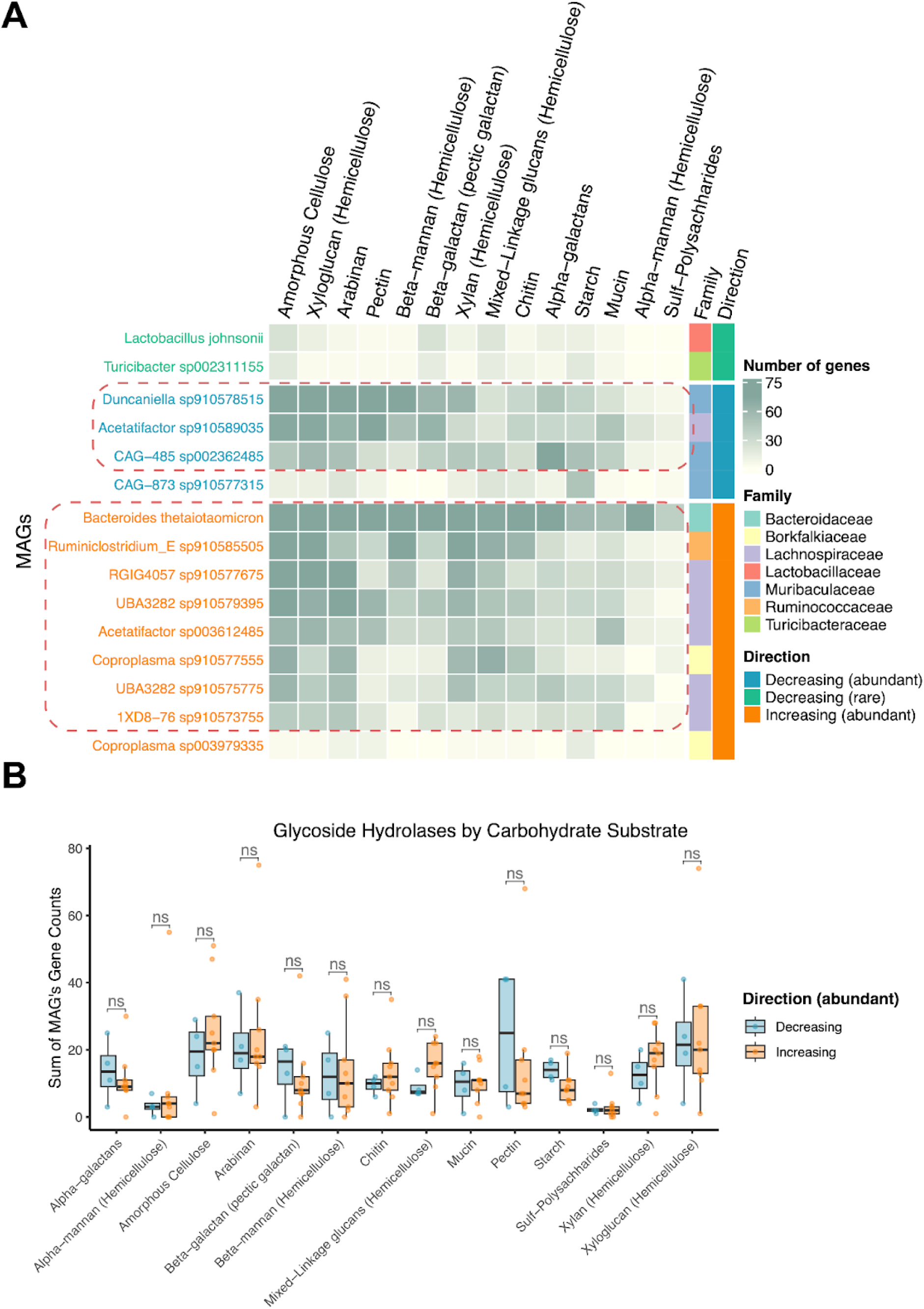
Carbohydrate substrate-resolved glycoside hydrolases (GHs) encoded by the most abundant changing MAGs and rare beneficial taxa. **(A)** A heatmap showing the number of GHs encoded by each MAG organized by carbohydrate substrate cleavage specificity according to DRAM’s annotations (from the Carbohydrate-Active enZymes Database or CAZy^70^). GHs that act on multiple substrates were accounted for in each substrate category separately. Increasing abundant taxa after spinal cord lesions are shown in orange, decreasing abundant taxa are shown in blue, and decreasing rare taxa are shown in green. Dotted boxes highlight MAGs enriched in GHs. (**B**) Boxplot comparisons of the number of GHs encoded by abundant increasing and decreasing MAGs organized by carbohydrate substrate cleavage specificity as defined in (A). For all boxplots, whiskers represent the 25th and 75th percentile ranges while *p*-values represent those from Wilcoxon tests, adjusted for multiple comparisons by the fdr (false discovery rate) method (see **Methods**). ns: *p* > 0.05.

**Fig. S7.**
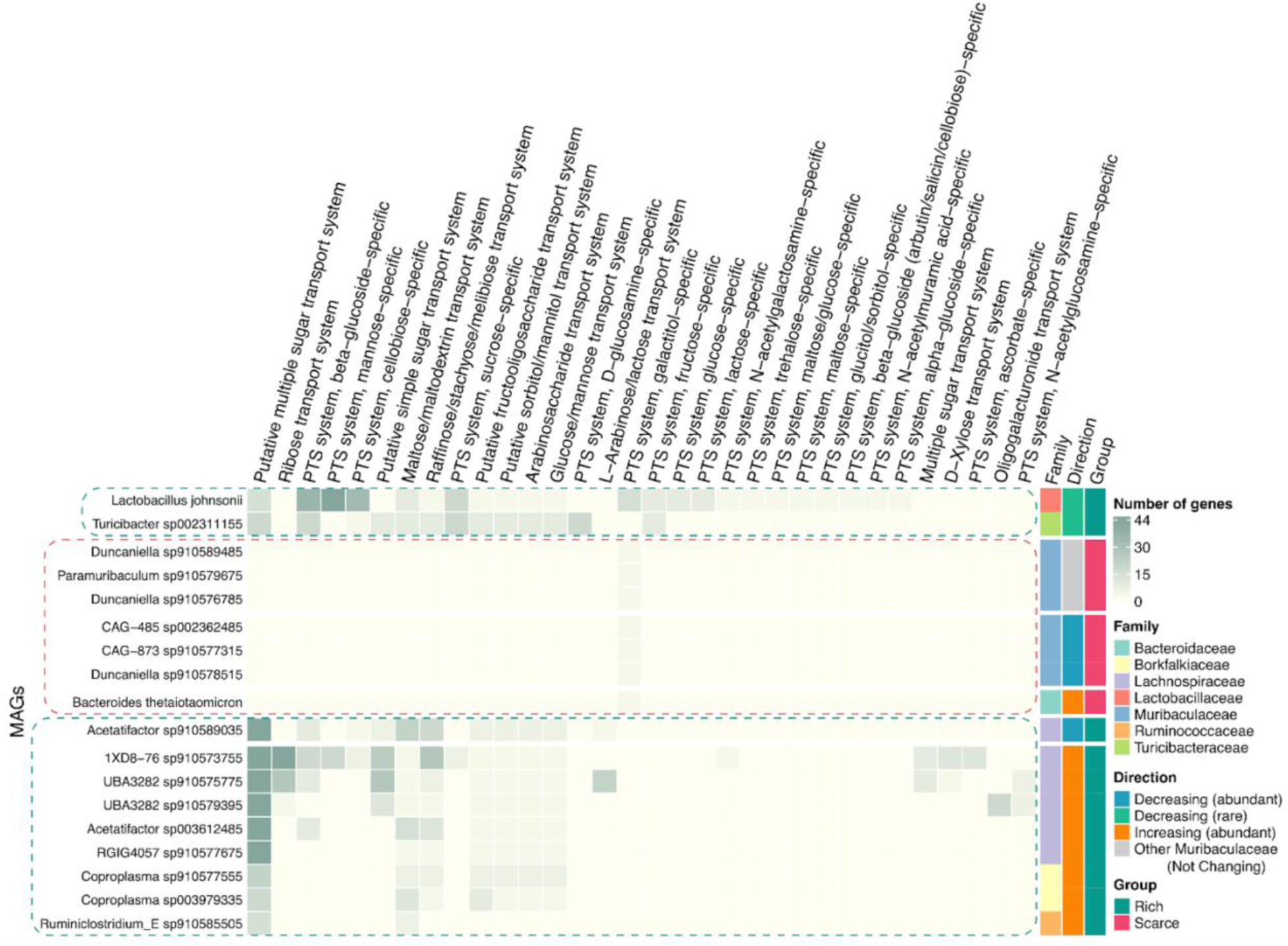
Sugar-resolved transporters and phosphotransferase systems encoded by the most abundant changing MAGs, other Muribaculaceae MAGs, and rare beneficial taxa. A heatmap showing the number of transporters and phosphotransferase systems (PTS) encoded by each MAG organized by sugar specificity according to DRAM’s annotations. Component II of PTS was used as a marker for the sugar-specific system by DRAM. Direction of change is indicated by orange for increasing abundant taxa after spinal cord lesions, blue for decreasing abundant taxa, grey for other non-changing Muribaculaceae MAGs, and green for decreasing rare taxa. All bacteriodales (highlighted with a red dotted box) scarcely encode sugar transporters and PTS, while rare beneficial taxa and abundant increasing taxa (both highlighted with green dotted boxes) are relatively rich in these sugar import systems.

**Fig. S8.**
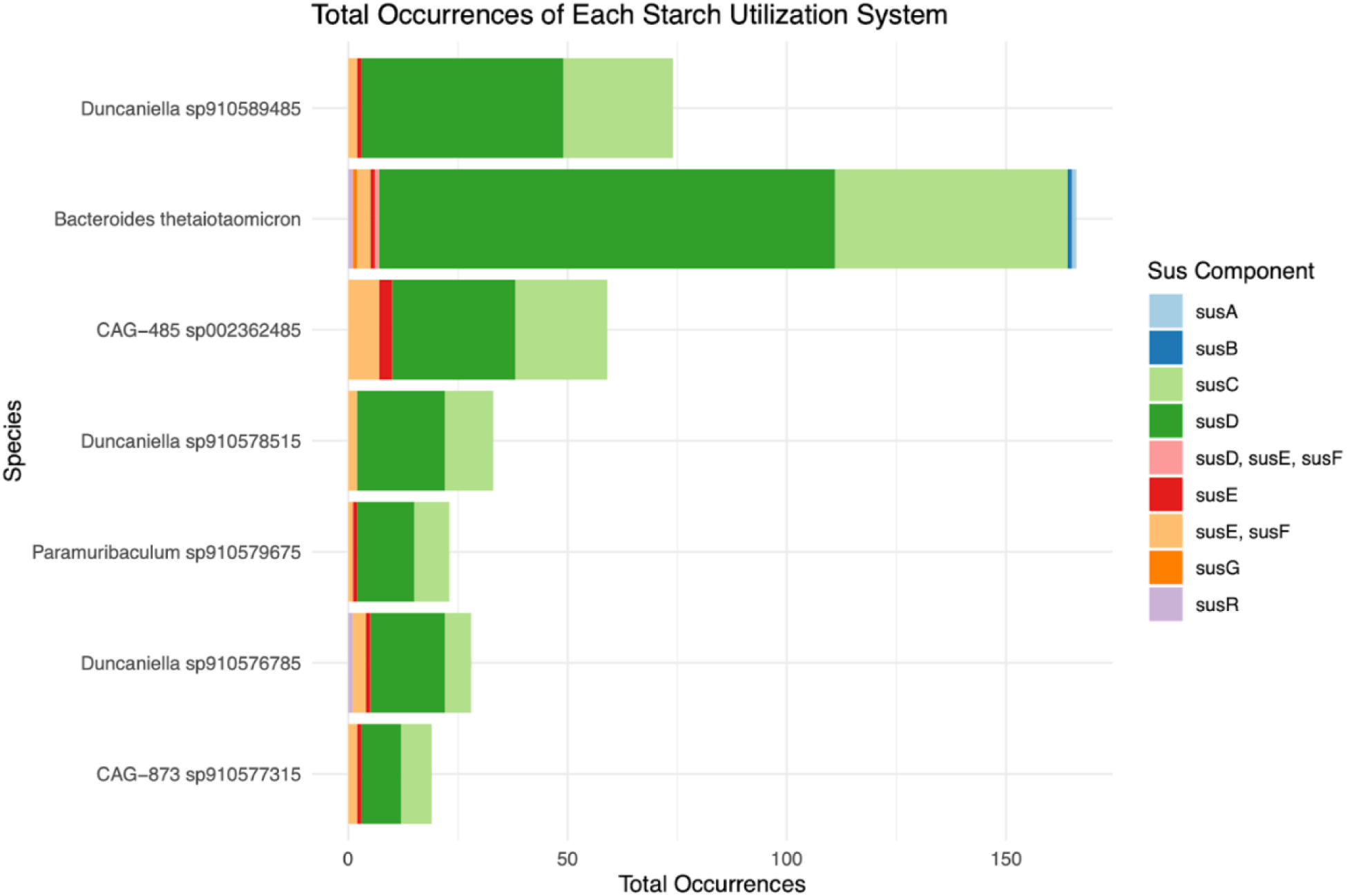
Prevalence of starch utilization system (SUS) in members of the bacteroidales order. Stacked bar chart showing the different SUS components detected in members of the bacteroidales order as defined in^65^ and annotated by DRAM. *B. thetaiotamicron*, the only increasing bacteroidales MAG after injury as seen in female mice data, encodes a markedly larger number of SUS homologs (using *susC* and *susD* as marker genes).

**Table S1.**
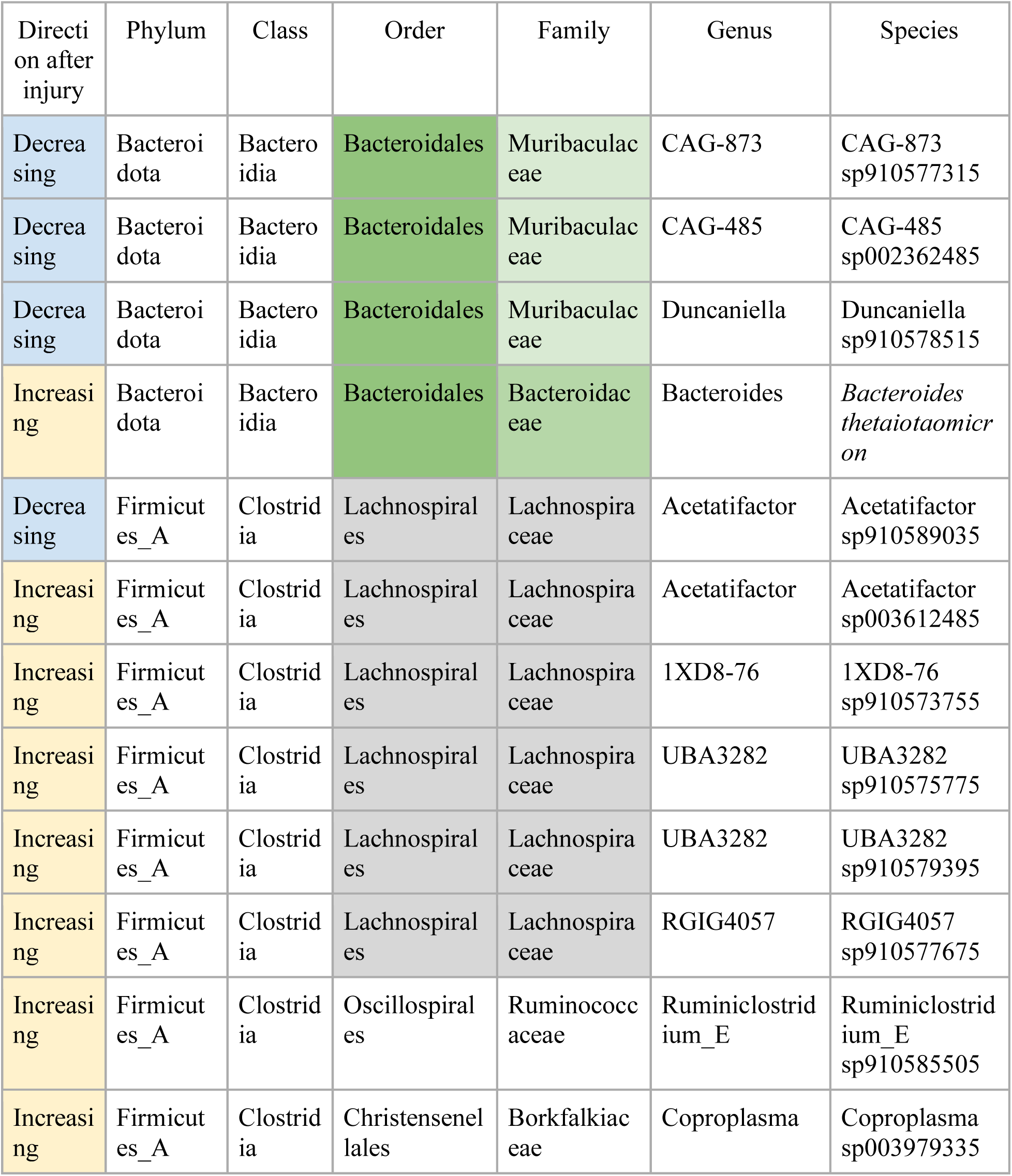
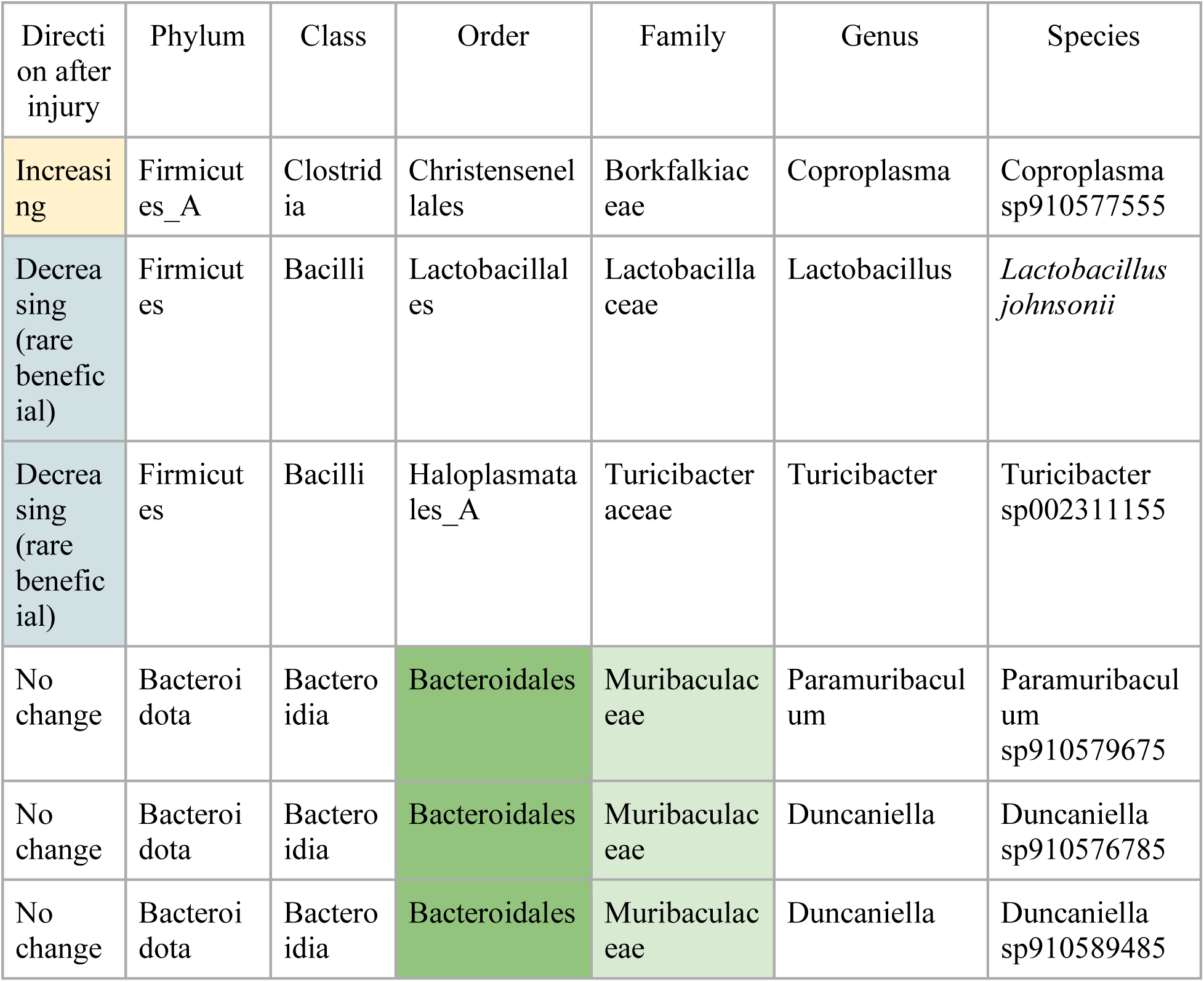
Taxonomic classification of the most abundant and rare beneficial changing MAGs, as well as non-changing Bacteroidales MAGs. Abundant decreasing and increasing MAGs are shown in blue and orange, respectively. Rare beneficial MAGs are shown in green, while non-changing MAGs are not colored. The Bacteroidales order is colored in green, with different shades showing different families (Muribaculaceae and Bacteroidaceae). Lachnospirales are colored in grey at the order and family (Lachnospiraceae) levels.

## Methods

### Animals and creating spinal cord lesions

All surgical and post-operative care procedures were performed in accordance with the Ohio State University Institutional Animal Care and Use Committee. Food (Teklad Global 14% Protein Rodent Maintenance Diet) and water were available at libitum. For metagenomic sequencing, N= 29 female C57BL/6 mice and N=30 male C57BL/6 mice from Jackson Laboratories (Bar Harbor, Maine) were used in this study. To prevent gut microbial cross-contamination due to co-habitation, all mice were single-housed upon arrival at our animal facility and for the duration of the study. Mice were anesthetized with an intraperitoneal cocktail of ketamine (80 mg/kg)/xylazine (10 mg/kg) after which a partial laminectomy was performed at the fourth thoracic spine (T4) or the tenth thoracic spine (T10). In mice modeling spinal cord injury (SCI), modified #5 Dumont forceps were inserted laterally and held closed for 3 seconds to produce a complete crush spinal cord injury. In total, 20 mice received a complete crush at T4, 20 mice received a complete crush at T10, and 19 mice received laminectomy (Lam) only (**Fig. 1A**).

Post-operatively, animals were hydrated with 2 mL Ringer’s solution (subcutaneous) for 5 days. Bladders were voided manually at least twice daily for the duration of the study. No prophylactic antibiotics were used during or after surgery. Fecal samples for metagenomic sequencing were collected pre-lesion, 1-week, 2-weeks, 3-weeks, 5-weeks, 9-weeks, and 6-months post-lesion. Mice were removed from their home cage and placed into a ventilated, aseptic polystyrene compartment and fresh fecal samples were collected from each mouse into sterile tubes and immediately frozen in liquid nitrogen. Mice were returned to their home cage after sample collection. At 6-months post-lesion, mice were anesthetized with ketamine/xylazine for tissue collection. Cecal contents were collected by removal of the cecum and extrusion of cecal contents into a sterile Eppendorf tube. All samples were immediately frozen in liquid nitrogen and stored at -80℃ until further use.

### Analysis of neurological function

Spinally-mediated neurological function was assessed using the Basso Mouse Scale for Locomotion (BMS score) at 6 months post lesion. Two experimenters assessed BMS scores for mice during a 4-minute interval in the open-field. Scores were tallied for each hind paw and then averaged to obtain a single score per animal. Lam mice have a score of 9 indicating normal locomotor function. Mice with spinal lesions have scores below 9, with scores varying as function of lesion level, with higher lesion levels creating greater neurological impairment across the body and correspondingly lower BMS scores. Deceased mice were assigned a score of 0 in this study.

Lactobacillus johnsonii isolation:

Fresh fecal samples were collected from naïve C57BL/6 mice (Jackson Laboratories) into sterile tubes and homogenized in 500ul sterile 0.1M PBS. 100ul of this homogenate was plated on agar plates made with De Man, Rogosa and Sharpe agar (Lactobacilli MRS agar; Neogen culture media NCM0035A), a selective culture medium designed to encourage growth of Lactobacillus species. These plates were cultured under anaerobic conditions using Mitsubishi™ AnaeroPack-Anaero Anaerobic Gas Generator (Thermo Scientific™ R681001) for 48hrs and numerous colonies were visible on the plates. To determine whether the resulting colonies were indeed L. johnsonii we selected 10 isolated colonies for further characterization. These colonies were expanded by culturing in MRS broth (MRS Broth; Millipore 69966) under static anaerobic conditions for 48hrs at 37C. After the incubation, the cultures were centrifuged to pellet the bacteria and cells were frozen in 40% glycerol-MRS broth and DNA was isolated from each culture using the Qiagen DNeasy PowerSoil Pro kit (Qiagen 47014). After DNA isolation, 16S PCR was run using universal 16S primers (GM3: AGA GTT TGA TCM TGG C; GM4: TAC CTT GTT ACG ACT T) following protocol (Natalie Solonenko 2016. 16S PCR Universal. protocols.io https://dx.doi.org/10.17504/protocols.io.c5my45). These primers amplify the 16S gene region for all bacteria (they are not genus or species specific). PCR products were then run on an agarose gel and gel-purified using Zymoclean Gel DNA Recovery Kit (Zymo Research; D4001). This gel-purified DNA was sent to the Genomics Core Lab at The Ohio State University for Sanger sequencing. The resulting sequences were then compared to all known DNA sequences using nucleotideBLAST (https://blast.ncbi.nlm.nih.gov/). The results of this query for isolate #10 are shown in Figure 1D. All results returned were specific for strains of *Lactobacillus johnsonii*, indicating that we have been successful in isolating and culturing this bacterium from the mouse gut microbiome. The same was true for all 10 isolates.

### Preparation of L. johnsonii glycerol stocks

After identifying that bacterial isolates were L. johnsonii, cultures were prepared to generate glycerol stocks for storing and future use in experimental studies. Frozen glycerol stocks (see above) were thawed and 1mL was transferred to a 50ml conical tube with 5mL sterile 0.1M PBS to remove glycerol. Cells were centrifuged at 4000xg for 10 minutes to pellet bacteria and supernatant was removed. Pellet was resuspended in 10mL of sterile MRS broth (MRS Broth; Millipore 69966) and cultured for 24hrs at 37C in anaerobic conditions (Mitsubishi™ AnaeroPack-Anaero Anaerobic Gas Generator; Thermo Scientific™ R681001). This 10mL culture was then divided evenly (3.3mL) into 3 new sterile 50mL conical tubes containing a final volume of 25mL. These cultures are incubated or 6-9hrs at 37oC in anaerobic chambers until absorbance/OD is between 0.35 and 0.55 when read at 600nm. We compared the OD values to a growth curve chart to determine approximate cfu/mL values. Bacteria are pelleted as above and resuspended in 40% glycerol MRS broth (MRS Broth; Millipore 69966) at a final concentration of 1 x 109 cfu/ml in cryovials and stored at -80C. To ensure accurate concentrations prior to use in animal studies, an aliquot if then thawed, washed and plated on MRS agar (Lactobacilli MRS agar; Neogen culture media NCM0035A) plates using serial dilutions to determine exact cfu/mL.

### Oral delivery of L. johnsonii

C57BL/6 female mice (Jackson Laboratories) received a mid-thoracic spinal cord lesion as described above. Mice received 1 x 108 cfu/mL L. johnsonii from stored glycerol stocks, washed in sterile 0.1M PBS, in 100ul sterile saline by oral gavage each day starting on day 1 post-injury through day 9 post-injury. Fecal samples were collected before and again 9 days post-lesioning. Fecal samples were homogenized in 500uL sterile 0.1M PBS and 100ul was plated on MRS agar (Lactobacilli MRS agar; Neogen culture media NCM0035A) to enumerate Lactobacillus colonies. From the same fecal samples, DNA was isolated using Qiagen DNeasy PowerSoil Pro kit (Qiagen 47014) for 16S sequencing (Novogene). Body weights were measured before surgery and throughout the 9-day survival period.

### Intestinal transit time assay

Total intestinal transit time was measured using a 6% Red Carmine dye assay following the protocol published in^36^. C57BL/6 female mice (Jackson laboratories) were given either a T9 complete crush spinal cord injury or a laminectomy (sham) control surgery (see surgical details above). Starting at 7 days post-injury, and weekly thereafter, mice were given 150uL of 6% red carmine (Sigma-Aldrich C1022) dissolved in a 0.5% methylcellulose (Sigma-Aldrich M0512) solution. After oral gavage of the 6% carmine solution mice were fasted for the duration of the assay. After 180 minutes, each mouse was placed in a clean cage and monitored at 5 minutes intervals to assess fecal pellets for the passage of the red dye (fecal pellets are visibly red color). The time of first red fecal pellet is recorded and whole gut transit time is calculated as the time interval between the gavage and the first observed red fecal pellet.

### DNA extraction and metagenomic sequencing

From 59 mice, 333 bulk metagenomic samples were prepared for sequencing (batch #1). Both fecal and cecum samples were extracted using a DNeasy PowerSoil Pro Kit (# 47014, QIAGEN) following the manufacturer’s instructions. The samples were then normalized to 12 ng/uL and sent to the JP Sulzberger Columbia Genome Center. At the center, using a Riptide kit from iGenomX for library preparation, sequencing was performed using Illumina NovaSeq 6000 targeting 26M 2×150bp reads per sample. Reads from bulk metagenomes ranged from 3.5 million paired-end reads to 64.5, with a median value of 25.1 million paired-end reads. For 11 samples with the lowest coverage, from 3.5 to 13.6 million reads, sequencing was redone at the same center, at 25M 2×100bp reads per sample. Library prep was performed using Nextera XT kit from Illumina (batch #2) at the Sullivan Lab before sending samples to the sequencing center. The prefix “Repeat_” was added to their sample labels in all metadata spreadsheets and codes. Reads ranged from 5.7 million paired-end reads to 26.8 for these 11 samples, with a median value of 24.2 million paired-end reads.

These samples starting with “Repeat_” were used in addition to the original ones for binning bacterial genomes but were discarded after calculations of abundances and further analyses due to having different read lengths (batch #2 having 2×100bp).

### Read quality control

Adapter trimming: Reads were trimmed with bbduk^71^ (https://jgi.doe.gov/data-and-tools/bbtools/) from the right end (ktrim=r) to keep ones longer than 30bp (minlength=30) by using a kmer of 23 (k=23) and allowing for a mismatch of 1 (hdist=1 hdist2=1) and a lower kmer at the ends of the read (mink=11). In addition, reads mapping to adapters, artifacts, phix and overrepresented sequences (i.e., low-complexity sequences) using “ref= adapters,artifacts,phix,<PATH-to- overrepresented_sequences>” to remove all reads having a 23-mer match to these commonly found sequences in Illumina runs.

Mouse reads removal: Reads mapping with a minimum identity of 95% to our mouse strain C57BL/6NJ (genome downloaded from the NCBI assembly database using the accession number “GCA_001632555.1”) were removed using bbmap^71^ (https://sourceforge.net/projects/bbmap). Additional settings were used in bbmap (maxindel=3 bwr=0.16 bw=12 quickmatch fast minhits=2 overwrite=t usemodulo).

Quality trimming: Next, using bbduk, reads with average quality of 10 or ones where base quality drops to 10 are trimmed from both sides keeping reads with a minimum length of 30 basepairs (qtrim=rl maq=10 maxns=0 minlength=30 trimq=10).

### Read-based profiling

Read-based profiling was performed using SingleM v 0.13.2 (https://github.com/wwood/singlem) using raw reads. First, the “pipe” command was run on raw reads and on the 263 set of MAGs. Operational taxonomic unit (OTU) tables resulting from each run (for paired reads of one sample) were then combined by sex. Finally, the “appraisal” method was used to assess how well these MAGs represented the whole community, resulting in **Fig. S1B**.

### Co-assembly and decontamination of scaffolds for binning (see Fig. S1A)

After quality control, all trimmed paired-end reads per mouse were combined and assembled using SPAdes (v3.14.1) in the meta mode (i.e. metaSPAdes^72^; https://github.com/ablab/spades). All scaffolds below 500bp were discarded. The scaffolds longer than 500bp were blasted (-task megablast) against an in-house contaminants database (see DOI: 10.5281/zenodo.13871704 with details in the ReadMe file). This database included the mouse genome (strain C57BL/6NJ) and genomes of non-gut microbes and viruses that were sequenced together with samples from this study (batches #1 and #2). Alignments that have 95% nucleotide query-subject identity covering 80% of the scaffold were considered contaminants and were therefore discarded.

After removing contaminants, trimmed reads from all bulk metagenomes were mapped to the set of co-assembled decontaminated scaffolds using CoverM v0.6.1 (https://github.com/wwood/CoverM) in the “contig” mode with the following parameters “--min-read-percent-identity .95 --min-read-aligned-percent .75 --discard-unmapped”. Resulting bam files were used as inputs for binning tools.

### Construction of MAGs, estimation of abundance, and taxonomic classification (see Fig. S1A)

Metabat2^73^ and Maxbin2^74^ (with the 107 marker gene set) were run with co-assemblies and per-sample bam files through UniteM v0.0.18 (https://github.com/dparks1134/UniteM) with a minimum contig length to be binned of 1500bp. The resulting bins from each tool were refined per mouse with metaWRAP v1.2 (bin_refinement module with the --keep-ambiguous flag, keeping bins with more than 50% completion). The quality of all bins was assessed with CheckM^75^ v 1.0.18. All 6635 bins from all 59 mice were then pooled with the 112 bins from^24^. The 6149 bins that were estimated to be more than 60% complete and less than 10% contaminated were de-replicated via Galah v0.3.1 (https://github.com/wwood/galah), resulting in 263 bins that captured redundant sequence variation resulting from mouse-constrained co-assembly. These 263 bins were used for further analysis.

Bins were taxonomically classified by the Genome Database Taxonomy Toolkit^76^ (GTDB-TK) v2.1.1 through the “classify_wf” module, which uses “release 207” of GTDB. The abundance of de-replicated MAGs was estimated using CoverM v0.6.1 (https://github.com/wwood/CoverM) in the “genome” mode with the trimmed mean method through the parameters “-m trimmed_mean --min-read-percent-identity .95 --min-read-aligned-percent .75 --discard-unmapped”.

### GTDB vs NCBI Taxonomy

Differences between NCBI and GTDB (release 207) include the following: (a) NCBI’s Firmicutes is divided into Firmicutes (includes order Bacilli), Firmicutes_A (includes order Clostridia), and Firmicutes_B (includes order Dehalobacteriia); (b) Many members of the families Bacteroidaceae, Prevotellaceae, Muribaculaceae are reclassified as Muribaculaceae, belonging to the phylum Bacteroidota (i.e, Bacteroidetes in NCBI); (c) Some members of the Clostridiaceae family (p Firmicutes; c Clostridia; o Eubacteriales) are classified as CAG-508 (p Firmicutes_A; c Clostridia; o TANB77) and include mostly unclassified species; members classified as family CAG-552 (p Firmicutes_A; c Clostridia; o Christensenellales) were largely unclassified firmicutes belonging to order Eubacteriales or novel orders; there have been also many events of reclassification in the class Clostridia from order Eubacteriales to Lachnospirales, Oscillospirales, Peptostreptococcales, and TANB77.

### Comparisons of genomic catalogs

First, genomes from different catalogs were collected. These include the Comprehensive Mouse Microbiota Genome Catalog (CMMG)’s 1573 species-level representative genomes (95% ANI) (https://ezmeta.unige.ch/CMMG/), the Integrated Mouse Gut Metagenomic Catalog (iMGMC)’s 1296 representative genomes (https://github.com/tillrobin/iMGMC), the Mouse Gastrointestinal Bacteria Catalogue (MGBC)’s 26,640 high-quality, non-redundant genomes (MGBC-hqnr_26640; https://github.com/BenBeresfordJones/MGBC), and CBAJDB’s 504 strain-level (99% ANI) medium-and high-quality genomes (CBAJDB_MQ-HQ; https://zenodo.org/records/8395759). Second, CheckM^75^ v1.0.18 was run on MAG catalogs to provide completeness and contamination information for each MAG – except for MGBC, for which MAG completeness and contamination data were extracted from genomes metadata (MGBC_md_26640). Third, for MGBC and CBAJDB (catalogs with no publicly available species-level representative genomes), Galah v0.3.1 (https://github.com/wwood/galah) was run (--ani 95 --precluster-ani 90) to get species-level representative genomes for MGBC (n=1091) and CBAJDB (n=113). Finally, at the strain level, MB6GC MAGs (n=499) were clustered with CBAJDB (n=504) using Galah (--ani 99 --precluster-ani 90) - **Fig. 1E**. At the species level, MB6GC MAGs (n=263) were clustered with CMMG, iMGMC, MGBC individually (**Fig. 1F**) using Galah (--ani 95 --precluster-ani 90). Additionally, MB6GC MAGs (n=263) were also clustered with MAGs from all representative genomes of the 4 catalogs together (**Fig. 1G**) using Galah (--ani 95 --precluster-ani 90). Stacked bar charts were plotted using ggplot2 (https://ggplot2.tidyverse.org) while heatmap (as in all heatmaps hereinafter unless otherwise stated) were made using the ComplexHeatmap package^77^ (https://github.com/jokergoo/ComplexHeatmap). All tables hereinafter were processed using the tidyverse package in R^78^.

### Alpha- and beta-diversity estimation

Beta-diversity was assessed using principal coordinate analyses (PCoAs) with bray-curtis dissimilarity matrices. Bray-curtis distances were calculated using the “vegdist” function from the R package “vegan” (https://github.com/vegandevs/vegan). Vegan’s “adonis2” function was used to perform permanova tests on bray-curtis distances with 9999 permutations. Longitudinal analyses were done by plotting bray-curtis distances to the mouse’s Day 0 sample to account for individual variations using ggplot2 (https://ggplot2.tidyverse.org). Statistical testing was performed via the wilcox_test function from the “rstatix” package (https://CRAN.R-project.org/package=rstatix), with the following parameters: “paired=FALSE, ref.group = ‘Lam’, detailed=TRUE, alternative=’less’, p.adjust.method =’fdr’. Significance was shown using the “ggpubr” package’s stat_pvalue_manual function (https://rpkgs.datanovia.com/ggpubr).

Alpha diversity was assessed by Shannon’s index using the “diversity” function from the vegan package. Shannon’s indices for samples were plotted by timepoint (i.e., samples at one time point) and as an aggregate (e.g., all samples post-injury). Longitudinal statistical comparisons shown in boxplots were made in a fashion similar to beta-diversity. Aggregate statistical comparisons shown in main figures were performed using ggpubr’s stat_compare_means, with the following parameters: method = “wilcox.test”, paired=FALSE, method.args = list(exact=FALSE), label = “p.signif”, hide.ns =TRUE, ref.group = “.all”.

### Data curation

Upon observing beta-diveristy in all samples after sham or spinal cord injury, samples were shown to cluster into 3 main groups. Since mice had their operations in 4 batches, at 4 different dates, we hypothesized that this might be one of the confounders. Colored by the day (i.e., date) on which the surgery was performed on mice, male mice having their operation performed at the first surgery day and female mice having theirs at the fourth day clustered closer to each other and differently from their respective larger male and female clusters (see **Fig. S1C**). These mice were excluded from further analyses (e.g., alpha and beta diversity, etc.). In addition, repeated samples (batch #2), having a different read length from batch #1, were also excluded from these analyses. Main male and female clusters were treated separately and as different cohorts.

### Statistical methods for determining differentially abundant taxa (Fig. 4, Fig. S3)

The trimmed-mean coverage values from CoverM were converted to relative abundance per sample after excluding repeated samples. Relative abundances were summed per each taxon and used for further tests. Considering the complex experimental design and the large number of taxa to be investigated (**Fig. 1**), it was challenging to test significance per time point due to losing statistical power, especially for female mice. Therefore, two statistical approaches were used to examine the data. First, samples collected from every two consecutive timepoints (e.g., “Day0 and 1week”, “1week and 3weeks”, and so on) were combined and tested for omnibus changes between the three experimental groups; the relative abundance values were fed into a Kruskal-Wallis test (from the kruskal.test function in R) followed by a Benjamini-Hochberg correction for multiple testing to check if there is a difference between this taxon’s abundance in Lam, T10, and T4 mice collectively. To examine the specific changes (e.g., differences between the taxon’s abundances in Lam vs T10 or Lam vs T4), statistically significant taxa from Kruskal-Wallis tests (adjusted *p*-values of less than 0.05) were then tested by a Wilcoxon signed-rank test (from the pairwise.wilcox.test function in R) followed by a Benjamini-Hochberg correction. No taxa turned out to be differentially abundant between Lam and T10 or T4 from the first timepoint pair (i.e., “Day0 and 1week”). Taxa deemed differentially abundant between injured and control (i.e., Lam) mice by these tests were then collected and plotted in **Fig. 4** and **Fig. S3**. Taxa were plotted depending on their abundance as follows: if the median abundance of the taxon (in any of the three treatments) was above 20% for phyla or 0.5% for species, it was plotted in the high-relative abundance plots. Second, these taxa were compared post-surgery (> Day0) with a Wilcoxon signed-rank test followed by fdr correction. Hence, consensus taxa plotted in **Fig. 4** and **Fig. S3** were both statistically significant upon testing the samples aggregated at the post-injury level (> Day 0) *and* at the two consecutive timepoints level.

### MAG functional annotation

DRAM v1.3.6 (https://github.com/WrightonLabCSU/DRAM) was used to annotate MAGs against multiple databases, including the Carbohydrate-Active enZymes Database or CAZy^70^. DRAM’s distillate outputs, *i.e.*, the “product.tsv” and the Carbon Utilization and Transporters tabs of the “metabolism_summary.xlsx,” were used to profile each MAG’s electron transport chain components (**Fig. S5**), GHs (**Fig. S6**), and sugar transporters (including PTS components; **Fig. S7**), respectively, whereas each MAG’s “Annotations” table was used to profile its PUL systems (**Fig. S8**). MAG-resolved heatmaps were generated for (i) the electron transport chain, (ii) GHs by counting the number of genes belonging to each “subheader” (representing a different carbohydrate substrate), including oligo- and backbone-cleavage enzymes, (iii) sugar transporters by counting the number of genes belonging to each “module” (representing a different transport system), and (iv) PUL systems by counting the number of genes whose annotations contain the strings “susA”, “susB”, “susC”,“susD”,“susE”, “susF”,“susG”, and “susR”.

### Temporal dynamics of select taxa

Relative abundances of select taxa were plotted longitudinally using ggplot2 (https://ggplot2.tidyverse.org). Statistical testing was performed via the wilcox_test function from the “rstatix” package (https://CRAN.R-project.org/package=rstatix), with the following parameters: “paired=FALSE, ref.group = ‘Lam’, detailed=TRUE, alternative=’two.sided’, p.adjust.method =’fdr’. Significance was shown using the “ggpubr” package’s stat_pvalue_manual function (https://rpkgs.datanovia.com/ggpubr).

## Data availability

All data produced in this study are publicly available. Curated data and all used scripts and databases can be found in the following Zenodo DOI: 10.5281/zenodo.13871704 (unpublished until review is complete). Raw reads are uploaded to NCBI SRA under the BioProject accession number: PRJNA1168861 (to download, see relevant tables in the Zenodo directory).

## Acknowledgements

We appreciate the help and support from all members of the Popovich and Sullivan laboratories, especially Natalie Solonenko, who participated in data sequencing. We also thank Dr. Darryl A. Wesener and Dr. Patrick H. Bradley, for valuable discussions and providing feedback. We also acknowledge the Center of Microbiome Science, Ohio Supercomputer Center, and the Riffomonas educational resource^79^. This work is funded by a National Institutes of Neurological Disorders and Stroke R35 award (grant no. 1R35NS111582) to P.G.P., The Belford Center for Spinal Cord Injury (P.G.P.), The Ray W. Poppleton Research endowment (P.G.P.), a Craig H. Nielsen Foundation Senior Research award (grant no. 890085) to K.A.K, and National Science Foundation awards ABI#2149505 and DBI#2022070 to M.B.S.

## Author Contributions

P.G.P., M.B.S., K.A.K., A.A.Z. conceived the project and experimental design. P.G.P., M.B.S., K.A.K., A.A.Z., and M.M. developed the data analysis plan. P.G.P., M.B.S., K.A.K., A.A.Z., M.M., G.J.S., and J.M.S. helped with data interpretation and writing of the manuscript. M.M. performed the statistical, profiling, functional, and bioinformatic analyses, and managed and coordinated responsibilities for the manuscript preparation. A.A.Z. guided and supervised statistical, profiling, functional, and bioinformatic analyses, performed the decontamination analyses, and drafted the manuscript with M.M. M.M., K.A.K., A.A.Z., and G.J.S. contributed to figure generation and functional analyses and, with P.G.P., contributed to improving illustrations. G.J.S. uploaded raw reads to SRA. K.A.K. performed the intestinal transit time assay, isolation and culture of *L. johnsonii, L. jonhsonii* supplementation experiments, murine spinal cord injuries, as well as daily animal care and sample collection. J.D. performed the DNA extraction for metagenomic sequencing. All authors read and reviewed the manuscript and approved it in its final form.

## Declaration of generative AI and AI-assisted technologies

During preparation of this work, Microsoft CoPilot and ChatGPT were used periodically to improve writing or shorten passages. After using it, the authors reviewed and edited the content as needed and take full responsibility for the content of the publication.

## References

1. Giampazolias, E. et al. Vitamin D regulates microbiome-dependent cancer immunity. Science 384, 428–437 (2024).

2. Wilde, J., Slack, E. & Foster, K. R. Host control of the microbiome: Mechanisms, evolution, and disease. Science 385, eadi3338 (2024).

3. Cryan, J. F. & Dinan, T. G. Mind-altering microorganisms: the impact of the gut microbiota on brain and behaviour. Nat Rev Neurosci 13, 701–712 (2012).

4. Foster, J. A. & McVey Neufeld, K.-A. Gut-brain axis: how the microbiome influences anxiety and depression. Trends Neurosci 36, 305–312 (2013).

5. Fülling, C., Dinan, T. G. & Cryan, J. F. Gut Microbe to Brain Signaling: What Happens in Vagus…. Neuron 101, 998–1002 (2019).

6. Bonaz, B., Bazin, T. & Pellissier, S. The Vagus Nerve at the Interface of the Microbiota-Gut-Brain Axis. Front Neurosci 12, 49 (2018).

7. Browning, K. N. & Travagli, R. A. Central nervous system control of gastrointestinal motility and secretion and modulation of gastrointestinal functions. Compr Physiol 4, 1339–1368 (2014).

8. Roager, H. M. et al. Colonic transit time is related to bacterial metabolism and mucosal turnover in the gut. Nat Microbiol 1, 16093 (2016).

9. Collins, S. M., Surette, M. & Bercik, P. The interplay between the intestinal microbiota and the brain. Nat Rev Microbiol 10, 735–742 (2012).

10. Renehan, W. E., Zhang, X., Beierwaltes, W. H. & Fogel, R. Neurons in the dorsal motor nucleus of the vagus may integrate vagal and spinal information from the GI tract. Am J Physiol 268, G780–790 (1995).

11. Hölzer, H. H. & Raybould, H. E. Vagal and splanchnic sensory pathways mediate inhibition of gastric motility induced by duodenal distension. Am J Physiol 262, G603–608 (1992).

12. Tanaka, S. et al. Vagus nerve stimulation activates two distinct neuroimmune circuits converging in the spleen to protect mice from kidney injury. Proc Natl Acad Sci U S A 118, e2021758118 (2021).

13. Deuchars, S. A. et al. Mechanisms underpinning sympathetic nervous activity and its modulation using transcutaneous vagus nerve stimulation. Exp Physiol 103, 326–331 (2018).

14. Bonaz, B., Sinniger, V. & Pellissier, S. The Vagus Nerve in the Neuro-Immune Axis: Implications in the Pathology of the Gastrointestinal Tract. Front Immunol 8, 1452 (2017).

15. Cryan, J. F., O’Riordan, K. J., Sandhu, K., Peterson, V. & Dinan, T. G. The gut microbiome in neurological disorders. Lancet Neurol 19, 179–194 (2020).

16. Valido, E. et al. Systematic review of the changes in the microbiome following spinal cord injury: animal and human evidence. Spinal Cord 60, 288–300 (2022).

17. Hou, K. et al. Microbiota in health and diseases. Signal Transduct Target Ther 7, 135 (2022).

18. Sorboni, S. G., Moghaddam, H. S., Jafarzadeh-Esfehani, R. & Soleimanpour, S. A Comprehensive Review on the Role of the Gut Microbiome in Human Neurological Disorders. Clin Microbiol Rev 35, e0033820 (2022).

19. Gohl, D. M. et al. Systematic improvement of amplicon marker gene methods for increased accuracy in microbiome studies. Nat Biotechnol 34, 942–949 (2016).

20. Bonk, F., Popp, D., Harms, H. & Centler, F. PCR-based quantification of taxa-specific abundances in microbial communities: Quantifying and avoiding common pitfalls. J Microbiol Methods 153, 139–147 (2018).

21. Arevalo, P., VanInsberghe, D., Elsherbini, J., Gore, J. & Polz, M. F. A Reverse Ecology Approach Based on a Biological Definition of Microbial Populations. Cell 178, 820–834.e14 (2019).

22. Kim, N. et al. Genome-resolved metagenomics: a game changer for microbiome medicine. Exp Mol Med 56, 1501–1512 (2024).

23. Eren, A. M. & Banfield, J. F. Modern microbiology: Embracing complexity through integration across scales. Cell 187, 5151–5170 (2024).

24. Du, J. et al. Spinal Cord Injury Changes the Structure and Functional Potential of Gut Bacterial and Viral Communities. mSystems 6, e01356–20.

25. Taylor, R. B. & Weaver, L. C. Spinal stimulation to locate preganglionic neurons controlling the kidney, spleen, or intestine. Am J Physiol 263, H1026–1033 (1992).

26. Mabon, P. J., LeVatte, M. A., Dekaban, G. A. & Weaver, L. C. Identification of sympathetic preganglionic neurons controlling the small intestine in hamsters using a recombinant herpes simplex virus type-1. Brain Res 753, 245–250 (1997).

27. Basso, D. M. et al. Basso Mouse Scale for locomotion detects differences in recovery after spinal cord injury in five common mouse strains. J Neurotrauma 23, 635–659 (2006).

28. Parks, D. H. et al. A standardized bacterial taxonomy based on genome phylogeny substantially revises the tree of life. Nat Biotechnol 36, 996–1004 (2018).

29. Leleiwi, I. et al. Exposing new taxonomic variation with inflammation - a murine model-specific genome database for gut microbiome researchers. Microbiome 11, 114 (2023).

30. Kieser, S., Zdobnov, E. M. & Trajkovski, M. Comprehensive mouse microbiota genome catalog reveals major difference to its human counterpart. PLoS Comput Biol 18, e1009947 (2022).

31. Beresford-Jones, B. S. et al. The Mouse Gastrointestinal Bacteria Catalogue enables translation between the mouse and human gut microbiotas via functional mapping. Cell Host Microbe 30, 124–138.e8 (2022).

32. Lesker, T. R. et al. An Integrated Metagenome Catalog Reveals New Insights into the Murine Gut Microbiome. Cell Rep 30, 2909–2922.e6 (2020).

33. Bryant, C. D. The blessings and curses of C57BL/6 substrains in mouse genetic studies. Ann N Y Acad Sci 1245, 31–33 (2011).

34. Solden, L. M. et al. Interspecies cross-feeding orchestrates carbon degradation in the rumen ecosystem. Nat Microbiol 3, 1274–1284 (2018).

35. Rodriguez, G. M. & Gater, D. R. Neurogenic Bowel and Management after Spinal Cord Injury: A Narrative Review. J Pers Med 12, 1141 (2022).

36. Koester, S. T., Li, N., Lachance, D. M. & Dey, N. Marker-based assays for studying gut transit in gnotobiotic and conventional mouse models. STAR Protoc 2, 100938 (2021).

37. Org, E. et al. Sex differences and hormonal effects on gut microbiota composition in mice. Gut Microbes 7, 313–322 (2016).

38. Groah, S. L. et al. Redefining Healthy Urine: A Cross-Sectional Exploratory Metagenomic Study of People With and Without Bladder Dysfunction. J Urol 196, 579–587 (2016).

39. Markle, J. G. M. et al. Sex differences in the gut microbiome drive hormone-dependent regulation of autoimmunity. Science 339, 1084–1088 (2013).

40. Yurkovetskiy, L. et al. Gender bias in autoimmunity is influenced by microbiota. Immunity 39, 400–412 (2013).

41. Rodríguez, A. E. et al. HIV medical providers’ perceptions of the use of antiretroviral therapy as nonoccupational postexposure prophylaxis in 2 major metropolitan areas. J Acquir Immune Defic Syndr 64 **Suppl 1**, S68–79 (2013).

42. Wu, Z. et al. Sex differences in colorectal cancer: with a focus on sex hormone-gut microbiome axis. Cell Commun Signal 22, 167 (2024).

43. Yoon, K. & Kim, N. Roles of Sex Hormones and Gender in the Gut Microbiota. J Neurogastroenterol Motil 27, 314–325 (2021).

44. Santos-Marcos, J. A., Mora-Ortiz, M., Tena-Sempere, M., Lopez-Miranda, J. & Camargo, A. Interaction between gut microbiota and sex hormones and their relation to sexual dimorphism in metabolic diseases. Biol Sex Differ 14, 4 (2023).

45. Razavi, A. C., Potts, K. S., Kelly, T. N. & Bazzano, L. A. Sex, gut microbiome, and cardiovascular disease risk. Biol Sex Differ 10, 29 (2019).

46. Saha, P. & Sisodia, S. S. Role of the gut microbiome in mediating sex-specific differences in the pathophysiology of Alzheimer’s disease. Neurotherapeutics 21, e00426 (2024).

47. Levenhagen, D. K. et al. Postexercise protein intake enhances whole-body and leg protein accretion in humans. Med Sci Sports Exerc 34, 828–837 (2002).

48. Gut dysbiosis impairs recovery after spinal cord injury | Journal of Experimental Medicine | Rockefeller University Press. https://rupress.org/jem/article/213/12/2603/42038/Gut-dysbiosis-impairs-recovery-after-spinal-cord.

49. Myers, S. A. et al. Following spinal cord injury, PDE4B drives an acute, local inflammatory response and a chronic, systemic response exacerbated by gut dysbiosis and endotoxemia. Neurobiology of Disease 124, 353–363 (2019).

50. Jing, Y. et al. Effect of fecal microbiota transplantation on neurological restoration in a spinal cord injury mouse model: involvement of brain-gut axis. Microbiome 9, 59 (2021).

51. Zhu, Y. et al. Exploration of the Muribaculaceae Family in the Gut Microbiota: Diversity, Metabolism, and Function. Nutrients 16, 2660 (2024).

52. Vacca, M. et al. The Controversial Role of Human Gut Lachnospiraceae. Microorganisms 8, 573 (2020).

53. Bai, Y. et al. Lactobacillus johnsonii enhances the gut barrier integrity via the interaction between GAPDH and the mouse tight junction protein JAM-2. Food Funct 13, 11021– 11033 (2022).

54. Jia, D.-J.-C. et al. Lactobacillus johnsonii alleviates colitis by TLR1/2-STAT3 mediated CD206+ macrophagesIL-10 activation. Gut Microbes 14, 2145843 (2022).

55. Lynch, J. B. et al. Gut microbiota Turicibacter strains differentially modify bile acids and host lipids. Nat Commun 14, 3669 (2023).

56. Harrigan, M. E. et al. Lesion level-dependent systemic muscle wasting after spinal cord injury is mediated by glucocorticoid signaling in mice. Sci Transl Med 15, eadh2156 (2023).

57. Zhu, J. et al. Probiotics and muscle health: the impact of Lactobacillus on sarcopenia through the gut-muscle axis. Front Microbiol 16, 1559119 (2025).

58. Chen, K. et al. Lactobacillus johnsonii CCFM1376 improves hypercholesterolemia in mice by regulating the composition of bile acids. Microbiome Res Rep 4, 6 (2025).

59. Fonseca, W. et al. Lactobacillus johnsonii supplementation attenuates respiratory viral infection via metabolic reprogramming and immune cell modulation. Mucosal Immunol 10, 1569–1580 (2017).

60. Xu, J. et al. Lactobacillus johnsonii Attenuates Liver Steatosis and Bile Acid Dysregulation in Parenteral Nutrition-Fed Rats. Metabolites 13, 1043 (2023).

61. Yang, G., Hong, E., Oh, S. & Kim, E. Non-Viable Lactobacillus johnsonii JNU3402 Protects against Diet-Induced Obesity. Foods 9, 1494 (2020).

62. Shaffer, M. et al. DRAM for distilling microbial metabolism to automate the curation of microbiome function. Nucleic Acids Res 48, 8883–8900 (2020).

63. Giuffrè, A., Borisov, V. B., Arese, M., Sarti, P. & Forte, E. Cytochrome bd oxidase and bacterial tolerance to oxidative and nitrosative stress. Biochim Biophys Acta 1837, 1178–1187 (2014).

64. Singh, R. P. Glycan utilisation system in Bacteroides and Bifidobacteria and their roles in gut stability and health. Appl Microbiol Biotechnol 103, 7287–7315 (2019).

65. Polysaccharide degradation by the Bacteroidetes: mechanisms and nomenclature - McKee - 2021 - Environmental Microbiology Reports - Wiley Online Library. https://enviromicro-journals.onlinelibrary.wiley.com/doi/10.1111/1758-2229.12980.

66. Grondin, J. M., Tamura, K., Déjean, G., Abbott, D. W. & Brumer, H. Polysaccharide Utilization Loci: Fueling Microbial Communities. Journal of Bacteriology 199, 10.1128/jb.00860-16 (2017).

67. Rakoff-Nahoum, S., Coyne, M. J. & Comstock, L. E. An ecological network of polysaccharide utilization among human intestinal symbionts. Curr Biol 24, 40–49 (2014).

68. Elhenawy, W., Debelyy, M. O. & Feldman, M. F. Preferential packing of acidic glycosidases and proteases into Bacteroides outer membrane vesicles. mBio 5, e00909–00914 (2014).

69. Takeuchi, T. et al. Gut microbial carbohydrate metabolism contributes to insulin resistance. Nature 621, 389–395 (2023).

70. Mori, S., Izumi, S. & Tomino, S. Complete nucleotide sequences of major plasma protein genes of Bombyx mori. Biochim Biophys Acta 1090, 129–132 (1991).

71. Bushnell, B., Rood, J. & Singer, E. BBMerge - Accurate paired shotgun read merging via overlap. PLoS One 12, e0185056 (2017).

72. Nurk, S., Meleshko, D., Korobeynikov, A. & Pevzner, P. A. metaSPAdes: a new versatile metagenomic assembler. Genome Res 27, 824–834 (2017).

73. Kang, D. D. et al. MetaBAT 2: an adaptive binning algorithm for robust and efficient genome reconstruction from metagenome assemblies. PeerJ 7, e7359 (2019).

74. Wu, Y.-W., Simmons, B. A. & Singer, S. W. MaxBin 2.0: an automated binning algorithm to recover genomes from multiple metagenomic datasets. Bioinformatics 32, 605–607 (2016).

75. Parks, D. H., Imelfort, M., Skennerton, C. T., Hugenholtz, P. & Tyson, G. W. CheckM: assessing the quality of microbial genomes recovered from isolates, single cells, and metagenomes. Genome Res 25, 1043–1055 (2015).

76. Chaumeil, P.-A., Mussig, A. J., Hugenholtz, P. & Parks, D. H. GTDB-Tk v2: memory friendly classification with the genome taxonomy database. Bioinformatics 38, 5315–5316 (2022).

77. Gu, Z. Complex heatmap visualization. Imeta 1, e43 (2022).

78. Wickham, H. et al. Welcome to the Tidyverse. Journal of Open Source Software 4, 1686 (2019).

79. Schloss, P. D. The Riffomonas YouTube Channel: An Educational Resource To Foster Reproducible Research Practices. Microbiol Resour Announc 12, e0131022 (2023).

